# Reinstatement of cortical outcome representations during higher-order learning

**DOI:** 10.1101/2020.05.28.121558

**Authors:** Lennart Luettgau, Emanuele Porcu, Claus Tempelmann, Gerhard Jocham

**Affiliations:** Biological Psychology of Decision Making, Institute of Experimental Psychology, Heinrich Heine University, Universitätsstrasse 1, 40225 Düsseldorf, Germany; Center for Behavioral Brain Sciences, Otto-von-Guericke University, Universitätsplatz 2, 39106 Magdeburg, Germany; Max Planck University College London Centre for Computational Psychiatry and Ageing Research, Russell Square House 10-12 Russell Square, WC1B 5EH London, UK; Department of Biological Psychology, Otto-von-Guericke University, Universitätsplatz 2, 39106 Magdeburg, Germany; Department of Neurology, Otto-von-Guericke University, Leipziger Strasse 44, 39120 Magdeburg

**Keywords:** Second-order conditioning, Learning, Decision Making, Orbitofrontal Cortex, Amygdala, fMRI

## Abstract

Naturalistic learning scenarios are characterized by infrequent experience of external feedback to guide behavior. Higher-order learning mechanisms like second-order conditioning (SOC) may allow stimuli that were never experienced together with reinforcement to acquire motivational value. Despite its explanatory potential for real-world learning, surprisingly little is known about the neural mechanism underlying such associative transfer of value in SOC. Here, we used multivariate cross-session, cross-modality searchlight classification on functional magnetic resonance imaging data obtained from humans during SOC. We show that visual first-order conditioned stimuli (CS) reinstate cortical patterns representing previously paired gustatory outcomes in the lateral orbitofrontal cortex (OFC). During SOC, this OFC region showed increased functional covariation with amygdala, where neural pattern similarity between second-order CS and outcomes increased from early to late stages of SOC. Our data suggest a mechanism by which motivational value is conferred to stimuli that were never paired with reinforcement.

## Introduction

Learning in naturalistic settings is characterized by infrequent direct encounters with rewarding or punishing stimuli (Gewirtz and Davis 2000). Hence, several stimuli or actions often need to be chained together, such that one stimulus serves as a proxy for another stimulus that might eventually predict reward (Gewirtz and Davis 2000). Exploiting such statistical regularities would allow agents to learn about the value of stimuli or actions that were never directly followed by reinforcement and support instrumental behavior in the absence of reinforcement or in new contexts. Second-order conditioning (SOC) is an example of higher-order learning that allows such associative transfer to stimuli that have never been directly paired with reinforcement (Pavlov 1927; Rizley and Rescorla 1972). In SOC, a first-order conditioned stimulus (CS_1_) is first paired with a motivationally salient event or stimulus (unconditioned stimulus, US). By virtue of this pairing, the CS_1_ is capable of evoking conditioned responses (CR, e.g. salivating, as in the presence of food), enabling it to function as conditioned reinforcer (Sharpe et al. 2017). Subsequently, another previously neutral conditioned stimulus (CS_2_) is paired with the CS_1_. Thereby, CS_2_ acquires incentive properties and is, like CS_1_, now able to elicit a CR. Despite its relevance for real-life learning phenomena (Gewirtz and Davis 2000; Parkes and Westbrook 2011) and decades of behavioral investigations (Rizley and Rescorla 1972; Barnet et al. 1991; Gewirtz and Davis 2000; Parkes and Westbrook 2011), surprisingly little is known about the neural mechanism underlying associative transfer of motivational value in SOC (Parkes and Westbrook 2011).

Building on models of memory reinstatement (Tonegawa et al. 2018), we hypothesized that during SOC, presentation of the CS_1_ would trigger reinstatement of the neural pattern representing the US with which it had previously been paired during first-order conditioning (FOC). This would allow linkage of CS_2_ and US representations by associative plasticity. Both amygdala (Hatfield et al. 1996; Gewirtz and Davis 1997, 2000; Setlow et al. 2002; Parkes and Westbrook 2011) and hippocampus (Gilboa et al. 2014), as well as orbitofrontal cortex (OFC) and ventral striatum (Mcdannald et al. 2013) have consistently been shown as the key structures involved in second-order learning and conditioned reinforcement (Hatfield et al. 1996; Gewirtz and Davis 1997, 2000; Setlow et al. 2002; Parkes and Westbrook 2011; Gilboa et al. 2014). Our analyses therefore focused on these regions.

To test our predictions, we combine a SOC paradigm and choice preference tests with cross-session (Stokes et al. 2009), cross-modality searchlight (Kriegeskorte et al. 2006) classification of functional magnetic resonance imaging (fMRI) data in healthy human participants. Participants first established Pavlovian associations between visual CS_1_ and appetitively or aversively valued gustatory US. During SOC, participants were exposed to associations between visual CS_2_ and previously learned CS_1_. In a subsequent preference test phase, participants were more likely to select directly (CS_1_) and indirectly (CS_2_) appetitively paired stimuli over aversively paired stimuli. These behavioral associative transfer learning effects were accompanied by CS_1_-related reinstatement of the neural patterns representing US in the lateral OFC and increased functional coupling between lateral OFC, amygdala/anterior hippocampus, and medial OFC during SOC. Furthermore, representations of second-order CS in the amygdala became more similar to US representations from early to late phases of second-order conditioning, indicating the acquisition of an association between CS_2_ and US patterns.

## Materials and Methods

### Participants

Participants were recruited from the local student community of the Otto-von-Guericke University Magdeburg, Germany by public advertisements and via online announcements. Only participants indicating no history of psychiatric or neurological disorder, no regular intake of medication known to interact with the central nervous system and participants not reporting quinine intolerance were included. Participants in both samples had normal or corrected-to-normal vision and did not report experience with Japanese kanjis or Chinese characters. All participants provided informed written consent before participation and received monetary compensation for taking part in the study. The study was approved by the local ethics committee at the medical faculty of the Otto-von-Guericke University Magdeburg, Germany (reference number: 101/15) and conducted in accordance with the Declaration of Helsinki. We assumed a medium effect size in a generic binomial test (effect size *g* = 0.25) for the main behavioral effects (choice probability > 0.50), for which a sample size of *N* = 23 would be necessary, given standard power (1–*β* = 0.80) and a one-tailed alpha-error probability of 0.05. We conducted two experimental studies. Since we used within-subject designs in both experiments and each participant experienced all conditions, effects of interest were tested within-subjects. There were no experimental groups and thus no randomization to experimental group was performed. 32 healthy adult volunteers (age: *M* = 24.16, *SD* = 3.61, range = 18 – 32 years, 15 males) participated in the fMRI study. We recruited 32 participants due to the fact that we expected ∼15% drop out in the fMRI study (due to compromised data, artifacts, falling asleep etc.). 20 healthy adult volunteers (age: *M* = 23.40, *SD* = 3.07, range = 19 – 30 years, 9 males), participated in the behavioral study. Participants self-reported high levels of education (50 of 52 subjects reported holding university entrance qualification degrees and received an average of 12.26 (*SD* = .95, range = 9 – 16) years of school education).

In the fMRI study, two participants were excluded from statistical analyses due to self-reports of having fallen asleep during the second-order conditioning scanning run. One additional participant had to be excluded due to a scanner malfunction and corruption of three of the five classifier training experiment scanning runs, thus leaving a total of *N* = 29 participants for final analyses.

In the fMRI study, volunteers participated in two sessions on two consecutive days (Figure 1A). The fMRI classifier training (see below) took place on the first day, and the learning experiment (next paragraph) was performed on the second day. In the behavioral study, participants attended one single session during which they performed the learning experiment.

**Figure 1.**
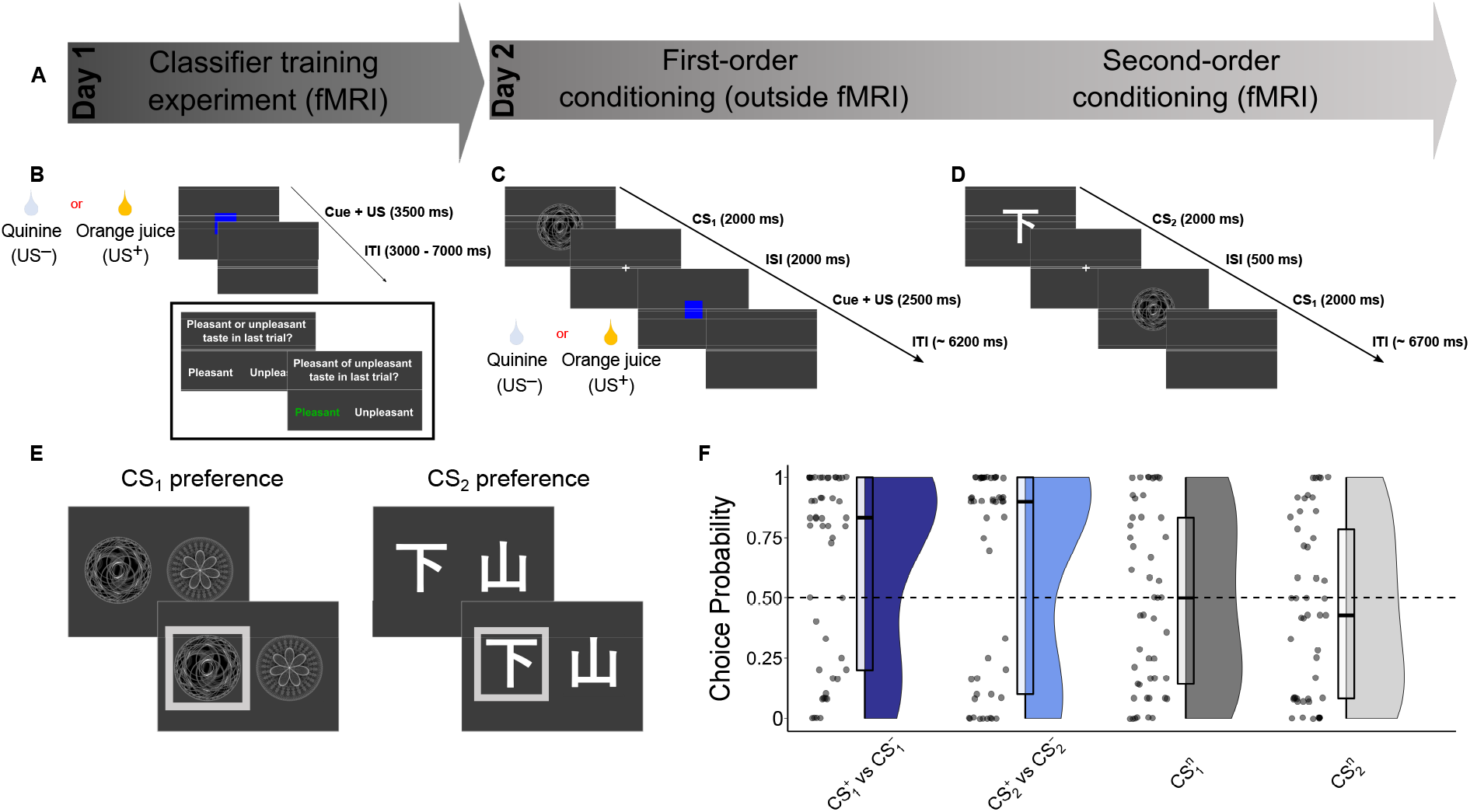
Experimental procedure, task schematic and behavioral results. A) Experimental procedure: Participants in the fMRI study (*N* = 29) performed both day 1 and day 2 (B-E), participants in the behavioral study (*N* = 20) only performed the procedures of day 2 (C-E, outside fMRI). B) Classifier training experiment: After rating of subjective value and intensity of both gustatory US (orange juice and quinine), in a total of five runs each US was administered twenty times (40 trials per run, 200 trials in total). Per trial, one US (1 ml bolus per trial) was delivered. In the fMRI study, the US bolus onset was preceded by a blue square. Trials were separated by an inter-trial-interval (ITI) marked by a grey screen. Participants performed a 0-back-style attentional control task. In 20% pseudo-randomly selected trials, participants were presented with probe trials in which they were asked to indicate which US they had received last (“pleasant” (US^+^) or “unpleasant” (US^−^)). C) First-order conditioning (outside MRI): In each trial, a CS_1_ was followed by an inter-stimulus interval marked by a fixation cross, and oral infusion of one US. Trials were separated by an ITI marked by a grey screen. CS_1_ were followed by a US with 80% probability. D) Second-order conditioning: In each trial, a CS_2_ was followed by an inter-stimulus interval marked by a fixation cross, and a CS_1_. CS_2_ were followed by a CS_1_ fully deterministically. Each trial was separated by an ITI marked by a grey screen. E) Choice preference test: Following SOC, participants were presented with two separate test phases consisting of repeated binary choices between pairs of CS_1_ (right) and pairs of CS_2_ (left) to assess behavioral signatures of first- and second-order conditioning. F) Behavioral results (combined across fMRI and behavioral study, *N* = 49): Raincloud plots (Allen et al. 2019) showing density of choice probability. Box plot center lines represent (pre-averaged) medians and box bottom/top edges show 25^th^/75^th^ percentile of the (pre-averaged) data, respectively.

### Learning experiment – ratings

Participants received written instructions for the experiment and were instructed once again on the computer screen. All experiments were programmed in MATLAB 2012b (v8.0.0.783, MATLAB and Statistics Toolbox Release 2012b, The MathWorks, Inc., Natick, MA, USA), using Psychophysics Toolbox (Brainard 1997) (version 3). Before and after the learning experiment, participants rated ten different round, greyscale fractal images (300 x 300 pixels) serving as first-order conditioned stimuli (CS_1_), ten white Japanese kanjis (Tamaoka et al. 2017) (250 x 250 pixels) serving as second-order conditioned stimuli (CS_2_) and gustatory stimuli as unconditioned stimuli (US). We used quinine-HCl (0.2 mmol/l solved in purified water) as aversive US and either chocolate milk (Nesquik, Nestlé, Switzerland) or orange juice (Milde Orange, EDEKA, Germany) as appetitive US (only orange juice in the fMRI study). Each participant received the same kind and amount of US per trial during the experiment. Ratings of subjective value/liking were assessed with the number buttons on a German (QWERTZ) computer keyboard from 1 (not liked) to 9 (very much liked). In the fMRI study, participants additionally rated gustatory stimuli regarding their subjective intensity levels from 1 (low intense) to 9 (very intense). Three fractals and kanjis rated closest to 5 (equivalent to “neutral”) were selected for first- and second order conditioning and their order was randomized before being associated with the US in first-order conditioning or with the first-order CS_1_ in second-order conditioning. To ensure motivational salience of the gustatory US, participants were instructed to abstain from food for 12 hours and on average reported having fasted for 13.28 (*SD* = 2.71) hours. Participants reported intermediate levels of hunger before the task on a paper-pencil visual analog scale (VAS), ranging from 0 (“not hungry”) to 100 (“very hungry”), *M* = 58.38 (*SD* = 29.61).

### Learning experiment – first-order conditioning

The first-order conditioning (FOC) phase of both the fMRI study and the behavioral study was performed outside of the scanner (Figure 1C). This was aimed at avoiding fatigue effects during the second-order conditioning phase, which was the phase of main interest to test our neural hypotheses. During FOC, participants were presented with CS^+^_1_ followed by the appetitive US^+^, and CS^−^_1_ followed by the aversive US^−^. CS_1_ were presented for 2000 ms, followed by an inter-stimulus interval (1000 ms) marked by a fixation cross, and oral infusion of one US (1 ml bolus per trial). Each CS_1_ was presented 50 times, amounting to 100 trials total. CS_1_ were followed by a US with 80% probability (40 trials of each CS_1_-US pair, 10 trials of CS_1_-no US per CS). US were delivered by a MATLAB code-controlled custom-made gustometer consisting of two high pressure single syringe pumps (AL-1000HP, World Precision Instruments, Saratoga, FL) operating 50 ml Luer lock syringes. Syringes were attached to Luer lock infusion lines (1,40 m length, 2 mm inner diameter) that participants held centrally in their mouths like drinking straws. In the fMRI study, infusion line position order (Q-O (*N* = 17) and O-Q (*N* = 15) for quinine (Q) and orange juice (O)), i.e. which US was delivered from the left or right infusion line, was counterbalanced across participants. In the fMRI study, the US bolus onset was preceded (500 ms) by a blue square that was presented for 2500 ms on the screen (Figure 1C). Participants were instructed to only swallow the US bolus upon offset of the blue square. Each trial was separated by an inter-trial-interval (ITI) marked by a grey screen. The ITI per trial was drawn from a discretized γ-distribution (shape = 6, scale = 1) truncated for an effective range of values between 3500 ms and 10,000 ms. Participants took self-paced breaks after each 10^th^ trial during which they could drink water. Importantly, the instructions did not contain information about the underlying associative structure of the experiment, aiming at leaving participants unaware of the associative learning process. Instead, participants were instructed to perform a simple attentional control task, during which they should respond as quickly and as correctly as possible by pressing the “y” button upon seeing the CS_1_ colored in red. Each CS_1_ was colored red in 10 % of the trials (90 % of trials grayscale image) and color did not predict US contingency. Performance during the task was rewarded with a bonus of 1 € (if > 70 % correct answers). Participants performed very well in the attentional control task (overall probability of correct answers: *M* = .99, *SD* = .05). In the fMRI study, both ratings and first-order conditioning were performed outside the MRI scanner.

### Learning experiment – second-order conditioning

In the fMRI study, second-order conditioning (SOC) was performed inside the MRI scanner. For SOC, participants were presented with CS_2_^+^ followed by CS_1_^+^, CS_2_^−^ followed by CS_1_^−^ and CS_2_^n^ followed by CS_1_^n^. CS_1_^n^ had not been presented during first-order conditioning and thus was not paired with any US. Since we assumed that both CS_2_^n^ and CS_1_^n^ should by design not elicit a neural representation of any of the US, CS_2_^n^ and CS_1_^n^ served as control stimuli. In each trial (Figure 1D), a CS_2_ (2000 ms) was followed by an inter-stimulus interval (500 ms) marked by a fixation cross, and a CS_1_ (2000 ms). CS_2_ were followed by a CS_1_ deterministically. Each trial was separated by an ITI marked by a grey screen. The ITI per trial was drawn from a discretized γ-distribution (shape = 7, scale = 1) truncated for an effective range of values between 4000 ms and 10,000 ms. Each CS_2_-CS_1_ pair was presented 50 times, amounting to 150 trials total. Again, instructions did not explicitly mention relational structures to be learned, but participants were instructed to perform a simple attentional control task. Participants were instructed to respond as quickly and as correctly as possible by pressing the “y” button (behavioral study) or with the right index finger on an MRI-compatible 4-button response box (fMRI study) upon seeing the CS_2_ tilted by a 45° angle. Each CS_2_ was tilted in 10 % of its presentations. Performance during the task was rewarded with a bonus of 1 € (if > 70 % correct answers). Participants performed very well in the attentional control task (overall probability of correct answers: *M* = .97, *SD* = .07).

### Learning experiment – choice preference test

Following SOC, participants were presented with two separate test phases consisting of repeated binary choices between pairs of CS_1_ and pairs of CS_2_ to assess behavioral signatures of first- and second-order conditioning. Participants were instructed to choose the stimulus they preferred (or “liked better”) on each choice trial, without being explicitly informed about the consequences of their choices. They chose between CS_1_^+^ and CS_1_^−^, and between CS_2_^+^ and CS_2_^−^, each choice being presented ten times (twelve times in the fMRI study). These decision trials were interleaved in pseudo-random order with fourteen (twelve in the fMRI study) choices between CS_1_^n^ or CS_2_^n^ and lure stimuli (fractal and kanjis, respectively) that had only been seen during pre-task rating. Neither CS_1_^+^, CS_2_^+^, nor CS_1_^−^, CS_2_^−^ were ever presented in comparison with CS_1_^n^, CS_2_^n^. We hypothesized preference for both CS_1_^+^ and CS_2_^+^ over CS_1_^−^ and CS_2_^−^, respectively, as expressed by choice probabilities above indifference criterion (choice probability > 0.5). The CS^n^-to-lure comparison trials were intended to rule out response biases related to mere exposure to the stimuli experienced during conditioning. We reasoned that if choices were reflecting acquired and transferred value and could not only be attributed to mere exposure-dependent response biases, choice probability in trials involving CS^n^-to-lure stimuli should not exceed the indifference criterion. Choice options were presented for 1500 ms on the right- and left-hand side of the screen. If participants did not respond within this time-window, a time-out message was displayed, and the respective trial was repeated at the end of the choice preference test. Order (left/right) of choice options was counterbalanced. Participants selected choice options by pressing the “y” (left option) or “m” (right option) button (behavioral study, left or right index finger on MR-compatible response box in the fMRI study). Participants were instructed to select the CS they preferred/liked more. Importantly, participants were never presented with the US related to their chosen or unchosen CS.

Following the choice preference test and after completing the behavioral paradigm, participants performed a post-experiment paper-pencil test. The test consisted of four questions assessing explicit knowledge about the associative structure of the second-order conditioning experiment. Participants were presented with a printed version of the ten fractal images and the ten kanji images that had initially been presented during ratings. Fractal and kanji images were numbered. We explicitly asked whether any of these kanjis (CS2) had been associated with a gustatory stimulus (US). Additionally, participants were asked which of the fractals (CS1) and which of the kanjis (CS2) presented on the printed page had also been presented during first- and second-order conditioning. Finally, we asked which fractals (CS1) and which kanjis (CS2) had been linked with which gustatory stimulus (US). Participants noted the respective numbers of the fractals and kanjis and linked these numbers with free-format words to describe the gustatory stimuli (e.g. “sweet” or “orange juice” to describe US+). We counted the number of correctly identified CS–US associations and calculated average (and standard deviation) to obtain a measure of explicit knowledge.

### Behavioral analyses

Data were analyzed using MATLAB 2019a (v9.6.0.1072779, The MathWorks, Inc., Natick, MA, USA) and RStudio (RStudioTeam 2019) (version 3.6.3, RStudio Team, Boston, MA) using custom analysis scripts. Choice probabilities for each CS in the fMRI and behavioral study were jointly analyzed. Each binary decision in which the respective CS was present (1 = selection of the respective target CS, 0 = selection of the alternative CS) for CS_1_^+^ and CS_2_^+^ (versus CS_1_^−^ and CS_2_^−^, respectively) and both first-order and second-order CS^n^-to-lure comparison trials was included in the models. We used six single-trial Bayesian multilevel generalized linear models (Equations 1–6), representing different hypotheses about which processes might have generated the choice data using the R package rethinking (McElreath 2020): 1) “Flat” model, modelling a single intercept parameter. This model assumes that choice behavior is invariant across subjects, studies and CS. 2) Model with CS-specific intercepts. This model assumes that choices vary only across CS but are invariant across subjects and studies. 3) Study-specific intercepts model. This model allows variation across studies (behavioral and fMRI study) but assumes invariance across subjects and CS. 4) Study-specific intercepts model with covarying CS-specific intercepts. This model includes variation across studies and CS and models covariation between study-specific and CS-specific effects. 5) Varying intercepts model with covarying CS-specific intercepts. This model allows subject-specific variation and includes covarying CS-specific effects. 6) “Full” model, combining individually varying intercepts and study-specific intercepts with covarying CS-specific intercepts. This model allows variation across subjects, studies and CS and also includes the covariation between these factors. We used Pareto-Smoothed Importance Sampling (PSIS) and the Widely Applicable Information Criterion (WAIC) for model comparisons and to find evidence for the best-fitting model. Within the best-fitting model, we hypothesized above-chance level choice probability for both CS_1_^+^ and CS_2_^+^ at the group-level. For both first-order and second-order CS^n^-to-lure comparison trials, we expected no difference from chance level for these choices. For hypothesis testing, we specified a region of practical equivalence (ROPE), i.e. an interval of parameter values (choice probability = [.45; .55]) representing the null hypothesis of no difference from chance level (“random”) choice behavior. Specifically, we expected that the 89% highest posterior density interval (HPDI) around the posterior parameter estimates – the interval containing 89% probability mass of the posterior distribution (i.e. the most credible posterior parameter values given the model and the data) – would not overlap with the ROPE for choices of CS_1_^+^ and CS_2_^+^ (versus CS_1_^−^ and CS_2_^−^, respectively). This expected non-overlap between HPDI and ROPE indicates that the most credible parameter values of the posterior probability distribution do not intersect with an a priori defined interval of parameter values that is indistinguishable from chance level (“random”) choice behavior. If there is no overlap between HPDI and ROPE, it is possible to reject the null hypothesis. However, we expected that the ROPE would overlap with the 89%-HPDI around the parameter estimates for both first-order and second-order CS^n^-to-lure comparison trials. This overlap would provide evidence for the null hypothesis, i.e, that the most credible parameter values are indistinguishable from chance level (“random”) choice behavior. Overlap between HPDIs and ROPEs was quantified using the R package bayestestR (Makowski et al. 2019).

Since we acquired binary choice data, we used a binomial distribution as likelihood function (e.g. Equation 1) and specified weakly informative prior and hyperprior probability distributions. Models were passed to RStan (Stan Development Team 2020) using the function “ulam” (rethinking). Using No-U-Turn samplers (NUTS; a variant of Hamiltonian Monte Carlo) in RStan and four independent Markov chains, we drew 4 x 10000 samples from posterior probability distributions (4 x 5000 warmup samples). Quality and reliability of the sampling process were evaluated with the Gelman-Rubin convergence diagnostic measure (*R̂* ≈ 1.00) and by visually inspecting Markov chain convergence using trace- and rank-plots. For all models fitted we found *R̂* = 1.00 for all parameters sampled from the posterior distribution. There were no divergent transitions between Markov chains for any of the models reported.

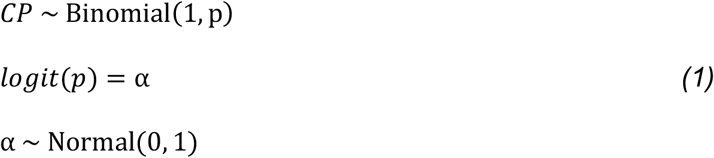

where *CP* is the binomially distributed choice probability for all CS (CS_1_^+^, CS_2_^+^, CS_1_^n^ or CS_2_^n^). *p* is the proportion of choices of CS.

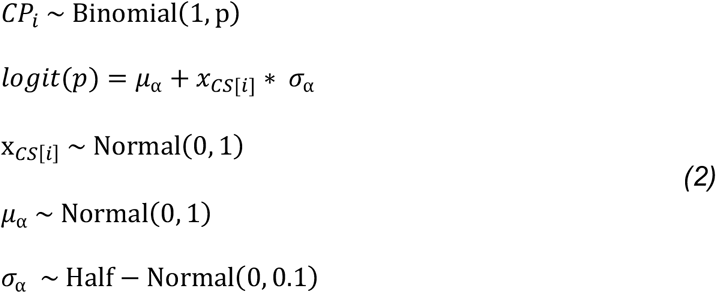

where *CP_i_* is the binomially distributed choice probability for each CS separately (CS_1_^+^, CS_2_^+^, CS_1_^n^ or CS_2_^n^). *p* is the proportion of CS choices. Note that the model is reparametrized to allow sampling from a standard normal posterior distribution.

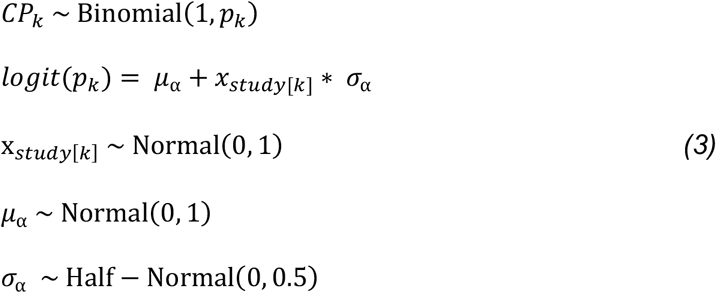

where *CP_k_* is the binomially distributed choice probability for the k-th study across all CS. *p_k_* is the proportion of CS choices. Again, the model is reparametrized to allow sampling from a standard normal posterior distribution.

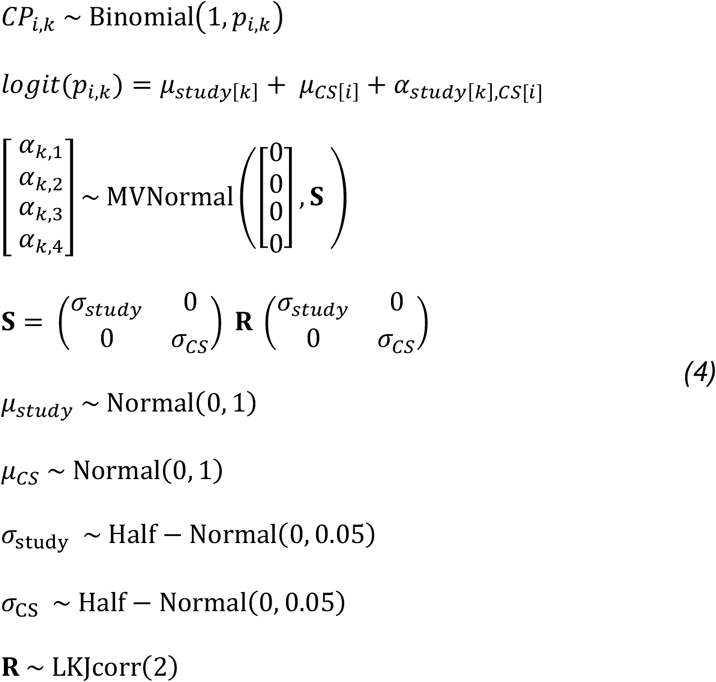

where *CP_i,k_* is the binomially distributed choice probability for the k-th study, covarying with the i-th CS-specific intercepts. *p_i,k_* is the proportion of CS choices.

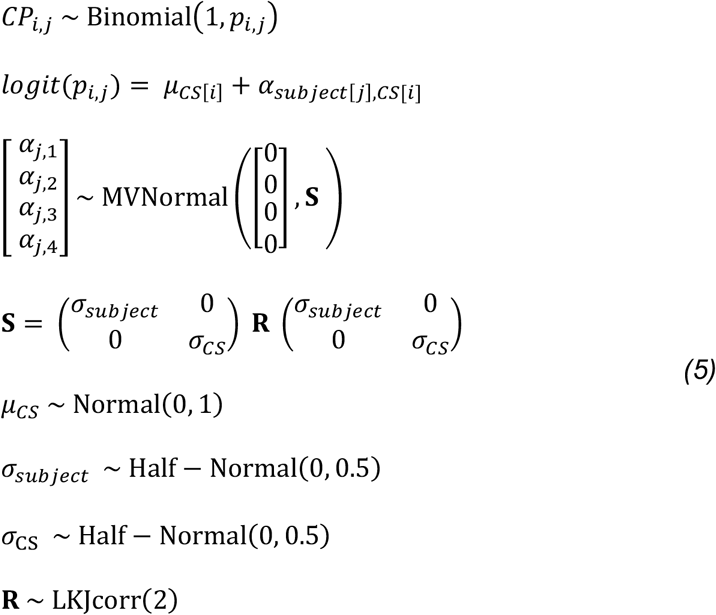

where *CP_i,j_* is the binomially distributed choice probability for the j-th subject, covarying with the i-th CS-specific intercepts. *p_i,j_* is the proportion of CS choices.

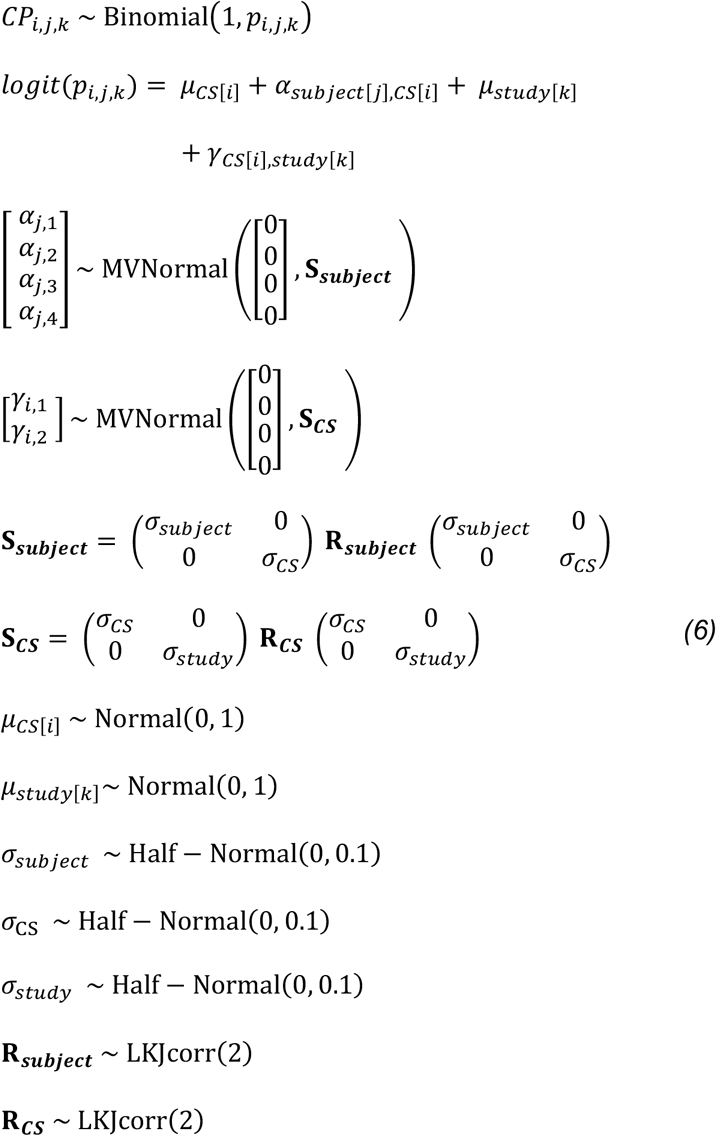

where *CP_i,j,k_* is the binomially distributed choice probability for the j-th subject covarying with the i-th CS-specific intercepts and the i-th CS-specific intercept covarying with the k-th study. *p_i,j,k_* is the proportion of CS choices.

### fMRI – classifier training experiment

The classifier training experiment was always performed on day 1 of two consecutive testing days. This was done to acquire unbiased estimates of the neural patterns representing the two US for training of a multivariate classification algorithm (multivariate pattern analysis, MVPA), before any association of the US with first- or second order CS had been acquired. Participants received oral instructions for the classifier training experiment and were instructed once again on the projection screen in the MRI. Before and after the experiment, participants rated gustatory US (aversive: quinine-HCL 0.1 mmol/l solved in purified water and appetitive: orange juice) according to subjective value/liking and intensity via button presses with their right and left index and middle fingers on a MRI-compatible 4-button response box. Ratings ranged from 1 (not liked/not intense) to 4 (very much liked/very intense) and were indicated by e.g. pressing the left middle finger corresponding to rating 1 or pressing the right index finger corresponding to rating 3. As on day 2, participants were instructed to abstain from food for 12 hours and on average reported to have fasted for 13.59 (*SD* = 2.18) hours. Participants reported intermediate levels of hunger before the experiment on a paper-pencil VAS, *M* = 48.50 (*SD* = 31.28). In each of the five total runs of the classifier training experiment, each US was administered twenty times (40 trials per run, 200 trials in total). Per trial, one US (1 ml bolus per trial) was delivered by a MATLAB code-controlled custom-made gustometer and Luer lock infusion lines (8.30 m length, 2 mm inner diameter). Participants held infusion lines centrally in their mouths like drinking straws. Additionally, infusion lines were fixed at absorbent materials attached to throat, chin and cheeks. The US bolus onset was preceded (500 ms) by a blue square that was presented in total for 4000 ms on the screen. Participants were instructed to only swallow the US bolus upon offset of the blue square. Each trial was separated by an ITI marked by a grey screen. The ITI was drawn from a uniform distribution with 5 discrete steps (range: 3000 – 7000 ms). Both US were presented in a pseudo-random order, thus reducing the influence of potentially confounding low-level features of the schedule (e.g. number of same/different US repetitions, different ITI lengths following each US). During each run of the classifier training experiment, participants performed a 0-back style attentional control task. After a pseudo-random 20% of trials, participants were presented with probe trials in which they were asked to indicate which US they had received last (“pleasant” (US^+^) or “unpleasant” (US^−^)) via button presses with their right and left index fingers on an MRI-compatible response box. Correct responses were rewarded with 0.05 € and incorrect responses or time-out trials (without a response by the participant within 2500 ms after onset of the probe trial) resulted in a 0.05 € penalty which would be summed up as a bonus upon completion of the experiment. On average, participants earned a bonus of 1.86 € (*SD* = .11) during the classifier training experiment. Performance during the 0-back attentional control task was generally high (overall probability of correct answers, excluding time-out trials: *M* = .93, *SD* = .05).

### fMRI – acquisition

During SOC, one run, and during the classifier training experiment, five runs of fMRI were acquired on a 3 Tesla Siemens PRISMA MR-system (Siemens, Erlangen, Germany), using a 64-channel head coil. Blood oxygenation level dependent (BOLD) signals were acquired using a multi-band accelerated T2*-weighted echo-planar imaging (EPI) sequence (multi-band acceleration factor 2, repetition time (TR) = 2000 ms, echo time (TE) = 30 ms, flip angle = 80°, field of view (FoV) = 220 mm, voxel size = 2.2 × 2.2 × 2.2 mm, no gap). Per volume, 66 slices covering the whole brain, tilted by approximately 15° in z-direction relative to the anterior– posterior commissure plane were acquired in interleaved order. The first 5 volumes of the functional imaging time series were automatically discarded to allow for T1 saturation. To ensure close spatial alignment of the slices acquired during both sessions, the AutoAlign technique provided by the vendor was applied. At the end of both testing days, a B_0_ magnitude and phase map was acquired to estimate field maps for B_0_ field unwarping during preprocessing (TR = 660 ms, TE 1 = 4.92 ms, TE 2 = 7.38 ms, flip angle = 60°, FoV = 220 mm). Additionally, before task-based fMRI on both days, a high-resolution three-dimensional T1-weighted anatomical scan (TR = 2500 ms, TE = 2.82 ms, FoV = 256 mm, flip angle = 7°, voxel size = 1 × 1 × 1 mm, 192 slices) covering the whole brain was obtained using a magnetization-prepared rapid acquisition gradient echo (MPRAGE) sequence. This scan was used as anatomical reference to the EPI data during the registration procedure. For all cross-session classification analyses, SOC EPI data was referenced to orientation of the classifier training experiment.

### fMRI – data preprocessing

All fMRI preprocessing steps were performed using tools from the Functional Magnetic Resonance Imaging of the Brain (FMRIB) Software Library (FSL, v5.0 and v6.0, Jenkinson et al., 2012). Preprocessing for each run of both classifier training task and SOC task included motion correction using rigid-body registration to the central volume of the functional time series (Jenkinson et al. 2002), correction for geometric distortions using the field maps and an n-dimensional phase-unwrapping algorithm (B_0_ unwarping, Jenkinson, 2003), slice timing correction using Hanning windowed sinc interpolation and high-pass filtering using a Gaussian-weighted lines filter of 1/100 Hz. EPI images were registered to the high-resolution structural image using affine linear registration (boundary-based registration) and then to standard (MNI) space using linear (12 degrees of freedom) and nonlinear registration (Andersson et al. 2007a, 2007b). Functional data was not spatially smoothed. We applied a conservative independent components analysis (ICA) to identify and remove obvious artefacts. Independent components were manually classified as signal or noise based on published guidelines (Griffanti et al. 2017). Before progressing to statistical analyses using general linear models (GLMs), the identified noise components were removed from the data by regressing components classified as noise from the functional imaging time series (using the FSL function fsl_regfilt).

### fMRI – searchlight classification analyses

We hypothesized that, during SOC, first-order CS would reinstate the neural pattern of the US with which they had previously been paired during FOC. We therefore expected that a multivariate classification algorithm trained on neural patterns evoked by the US (during the classifier training task) would be able to correctly predict the class of a paired CS_1_ during SOC. Hence, we used a cross-session (Stokes et al. 2009), cross-modality searchlight (Kriegeskorte et al. 2006) classification approach to identify brain regions in which the CS class during SOC could be predicted based on training the classifier during the classifier training experiment on day 1. GLMs were fitted into pre-whitened data space to account for local autocorrelations (Woolrich et al. 2001). For the searchlight classification analyses, the individual level (first level) GLM design matrix per run and participant of the classifier training experiment included four box-car regressors in total. Two regressors coded for onsets and durations of both US (each modelled as single events of 4000 ms duration) and two regressors coded onsets and durations of left and right button presses (delta stick functions on the recorded time of response button presses) and the six volume-to-volume motion parameters from motion correction during preprocessing were entered to account for residual head motion. The SOC run per participant included five box-car regressors in total. Three regressors coding for onsets and durations of all combinations of CS_2_-CS_1_ (CS_2_^+^/CS_1_^+^, CS_2_^−^/CS_1_^−^, CS_2_^n^/CS_1_^n^) trials (each modelled as single events of 4500 ms duration, due to the 100 % contingency of CS-CS pairs), one regressor coding onsets and durations of right button presses (delta stick functions on the recorded time of response button presses), one regressor coding onset and duration of the within-run pause (45 sec), and the six volume-to-volume motion parameters from motion correction during preprocessing were entered. Please note that for the SOC analyses, the CS_2_-CS_1_ presentations were modeled as single events. The close temporal succession of the two stimuli with a fixed inter-stimulus interval was chosen to obtain robust conditioning effects. This makes it difficult to obtain estimates of neural activity that can be unambiguously attributed to either the first- or second-order stimulus. Therefore, for this analysis (but see below for neural pattern similarity analyses) we chose to model them as single events. Please note however that this analysis is not aimed at separating out effects specific to either CS_1_ or CS_2_. Instead, we aim to detect reinstatement of US representations. If such reinstated US representations can be detected, they indeed can only be faithfully attributed to the CS_1_, as the CS_2_ had no association with the US.

Additionally, we performed an exploratory multivariate searchlight classification to distinguish between the spatial activation patterns (*t* maps) of US^+^ versus US^−^ across the whole brain during the classifier training experiment. The aim of this analysis was to present an overview over the whole-brain patterns allowing classification of the US. This analysis resulted in one whole-brain map of classification accuracies per participant. The classification analysis was performed using a 5-fold cross-validation scheme, iteratively training the classification algorithm on the difference between US^+^ versus US^−^ on 4 (out of 5 runs) labeled training data sets and then assessing the out-of-sample predictive accuracy of the classifier on the remaining, left-out data set (1 out of 5 runs).

Regressors were convolved with a hemodynamic response function (γ-function, mean lag = 6 s, SD = 3 s). Each first level GLM included two or three (for classifier training experiment or SOC, respectively) contrasts to estimate individual per run *t*-statistic maps for each US or each CS_2_-CS_1_ pair (for classifier training experiment or SOC, respectively). For SOC, the activation pattern of CS_2_^n^/CS_1_^n^ was subtracted from CS_2_^+^/CS_1_^+^ and CS_2_^−^/CS_1_^−^ activation patterns. This was intended to account for general activation expected to result from visually presented CS and the visually presented swallowing cue coinciding with US presentation, which could have confounded classification accuracies.

All classification-based analyses were conducted in subject native (fMRI) space. Per-participant *t*-statistic maps were subjected to linear support vector machine (C-SVM, cost parameter C = 1) classification using the MATLAB-based multivariate pattern analysis toolbox CoSMoMVPA (Oosterhof et al. 2016). For cross-session, cross-modality classification, a classifier was trained on the spatial activation patterns of US^+^ versus US^−^ (obtained from classifier training) and tested on the patterns of CS_2_^+^/CS_1_^+^ versus CS_2_^−^/CS_1_^−^ during SOC within 3-mm searchlight spheres across the whole brain. For the exploratory multivariate searchlight classification between US^+^ versus US^−^ patterns, we similarly used a 3-mm searchlight sphere across the whole brain. Each searchlight sphere classification accuracy was mapped to the center voxel of the sphere, resulting in one whole-brain map of classification accuracies per participant. Additionally, in cross-session, cross-modality classification we repeated this procedure 100 times per participant with randomly permuted class labels in the training data set (US^+^ versus US^−^) to create chance level maps. Before group level statistics, the normalization parameters obtained from preprocessing were applied to both whole-brain classification accuracy maps and chance level maps for normalization to MNI space. The resulting normalized whole-brain maps were spatially smoothed with a Gaussian kernel (5 mm full-width half-maximum).

For cross-session, cross-modality classification, we used small-volume correction (*P_SVC_*) to assess significance of clusters with above-chance level (0.5) classification accuracy. To correct for multiple comparisons, we used a meta-analysis-based explicit mask for functional activations related to the term “taste” in the Neurosynth data base (Yarkoni et al. 2011) (www.neurosynth.org), thresholded at *z* > 7. This value was chosen such that only meta-analytic clusters of activation, but not single voxels, survived thresholding. The Neurosynth-based mask encompassed, aside from the lateral OFC, functional activation clusters in bilateral anterior insula. In addition, we evaluated an anatomically and functionally defined region-of-interest (ROI) for the lateral orbitofrontal cortex (lOFC), our a priori ROI. This region has been implicated in higher-order gustatory processing, such as motivational aspects and discrimination of taste (Small et al. 1999; Kobayashi et al. 2004; Miranda 2012), but also representation and adaptive changes of stimulus-outcome associations (Klein-Flügge et al. 2013; Boorman et al. 2016; Jocham et al. 2016; Luettgau et al. 2020). We thus reasoned that this brain region would be a well-suited candidate for associative coupling between visual conditioned stimuli and gustatory outcomes and the proposed associative transfer learning. We used two different approaches for lOFC ROI definition: 1) an independent anatomical mask of the lateral OFC (Harvard-Oxford Atlas) and 2) an independent functional ROI from a gustatory mapping study by Benz and colleagues (K. Benz, personal communication, 12/2019). This approach aimed at reducing inferential limitations related to arbitrary ROI definition.

Within the meta-analysis-based mask and the ROIs, we computed group-level random-effect cluster-statistics corrected for multiple comparisons in small volumes as implemented in CoSMoMVPA (Oosterhof et al. 2016). In brief, for 50,000 iterations, we randomly selected one chance level map per participant, calculated a group level z-statistic map for the respective permutation and finally compared the resulting cluster sizes drawn from this empirical null distribution (50,000 samples) to the clusters in the “real” classification accuracy map (Stelzer et al. 2013). Since classifications accuracies below chance level are generally limited in interpretability, we considered clusters significant if *P_SVC_* < .05 (one-tailed). Please note that due to the exploratory nature and illustrative purpose of the exploratory multivariate searchlight classification between US^+^ versus US^−^ patterns, no multiple comparisons correction using random-effects cluster-statistics was performed in this analysis.

Additionally, we split participants into a high- or low-bias group, depending on their preference for the CS_2_^+^ (vs. CS_2_^−^) and compared classification accuracies within the small-volume corrected cluster. The cluster ROI was built in MNI space and the ROI was then back-projected into subject native space using inverse normalization parameters obtained during preprocessing to extract individual averaged classification accuracies from the ROI. Both groups’ average extracted classification accuracies were separately tested against chance level (0.5) using one-sample Wilcoxon signed-rank tests. We report measures of effect size Cohen’s *U3_1_* for one-sample Wilcoxon signed-rank tests (range: 0 – 1, .5 indicating no effect), calculated in the MATLAB-based Measures-of-Effect-Size-toolbox (Hentschke 2020).

### fMRI – functional covariation analyses

Our cross-session, cross-modality searchlight classification during SOC identified a cluster in the lOFC (see Results). We investigated functional covariation between lOFC and the rest of the brain during SOC using this lOFC cluster as a seed region in a psychophysiological interaction (PPI) analysis. For each participant, we set up two separate first-level PPI GLMs. The first PPI GLM was aimed at investigating general differences in BOLD signal covariation for both CS_2_^+^/CS_1_^+^ and CS_2_^−^/CS_1_^−^ compared with the control stimuli that had not been paired with a US, CS_2_^n^/CS_1_^n^. It contained the following regressors: 1) the BOLD timeseries (preprocessed functional time series) of the lOFC seed as physiological regressor, 2) onsets and durations of CS_2_^+^/CS_1_^+^ and CS_2_^−^/CS_1_^−^ pairs (magnitude coded as 1) and CS_2_^n^/CS_1_^n^ (magnitude coded as –1) as psychological regressor and 3) the interaction between 1) and 2) as psychophysiological interaction regressor. The psychological regressor was convolved with a hemodynamic response function (γ-function, mean lag = 6 s, *SD* = 3 s). The second PPI GLM aimed at investigating specific differences in covariation between CS_2_^+^/CS_1_^+^ and CS_2_^−^ /CS_1_^−^. It contained the following regressors: 1) the BOLD timeseries (preprocessed functional time series) of the lOFC seed as physiological regressor, 2) onsets and durations of CS_2_^+^/CS_1_^+^ (magnitude coded as 1) and CS_2_^−^/CS_1_^−^ pairs (magnitude coded as –1) as psychological regressor and 3) the interaction between 1) and 2) as psychophysiological interaction regressor. The psychological regressor was convolved with a hemodynamic response function (γ-function, mean lag = 6 s, *SD* = 3 s).

In addition to the main effects of the PPI regressors, we also investigated whether task-related functional covariation of lOFC with the rest of the brain was related to our behavioral index of second-order conditioning (choice probability of CS_2_^+^ vs. CS_2_^−^). For this, we entered the averaged second-order choice preference test data per subject as an additional regressor to the group level GLMs. In all analyses, contrast images from the first level were then taken to the group level and analyzed using mixed-effects analyses in FLAME1+2 (Beckmann et al. 2003). We used cluster-based correction with an activation threshold of Z > 2.3 and a cluster-extent threshold of *P* < .05 at whole-brain level to control for multiple comparisons.

### fMRI – neural pattern similarity analyses

In addition to our classification-based approach, we investigated whether there was evidence for changes of neural pattern similarity across second-order conditioning. Specifically, we used a least-squares separate (LS-S) approach (Mumford et al. 2012) to deconvolve single-trial estimates of neural patterns representing CS_2_ and CS_1_. We then used these patterns to perform a template-based neural pattern similarity analysis (Wimber et al. 2015) (a variant of representational similarity analysis, RSA (Kriegeskorte et al. 2008)) between patterns of the two US during the classifier training experiment and all trial-specific CS_2_/CS_1_ patterns during the SOC run. We estimated two subject-specific GLMs per trial, one for both CS_1_ and CS_2_, containing the following regressors: 1) onset and duration of the respective trial to be estimated (each modelled as a single event of 2000 ms duration), 2) onsets and durations of the remaining CS_1_ or CS_2_ trials, respectively, 3) onsets and durations of all other CS_1_ or CS_2_ trials, respectively. Additionally, one regressor coding onsets and durations of right button presses (delta stick functions on the recorded time of response button presses), one regressor coding onset and duration of the within-run pause (45 sec), and the six volume-to-volume motion parameters from motion correction during preprocessing were entered. For example, the GLM of the 25^th^ trial of CS_1_^+^, contained one regressor (onset and duration) modelling this particular trial and one separate regressor (onsets and durations) for all remaining CS_1_ trials but the 25^th^ trial of CS_1_^+^ (149 other trials in total) and one separate regressor (onsets and durations) modelling all CS_2_ trials. This procedure is assumed to allow for better deconvolution of events happening within close temporal proximity than standard least-squares-based approaches, which generally do not distinguish between overall and trial-specific error terms (Mumford et al. 2012; Abdulrahman and Henson 2016). As we have detailed above (paragraph “fMRI – searchlight classification analyses”), the temporal relation between CS_1_ and CS_2_ makes it inherently difficulty to estimate separate activation patterns. While, for the searchlight classification, it is not important to separate out the individual CS contributions to the neural pattern, our analyses on changes in neural pattern similarity across the SOC phase do make different predictions for CS_1_ and CS_2_. We therefore used the deconvolution approach described above to mitigate the issue, but caution is warranted in interpreting these results.

Each first level GLM included one contrast to model activation related to the respective trial of interest versus baseline. The a priori ROIs of the bilateral amygdala were built in MNI space and back-projected into subject native space using inverse normalization parameters obtained during preprocessing. We used these individual ROIs for spatially constrained multivoxel pattern extraction from the respective contrast *t*-value maps. Similarity-based analyses were carried out using CoSMoMVPA (Oosterhof et al. 2016). We used 1−Pearson’s product-moment correlation coefficient (1−*r*) as a measure of pairwise dissimilarity between trial-specific neural patterns and the US template. Within-subject pairwise neural dissimilarity was subtracted from 1 (to create a measure of neural pattern similarity) and Fisher-*Z* transformed to approximate normally distributed data more closely. CS_1_^n^-US^−^/CS_2_^n^-US^−^ and CS_1_^n^-US^+^/CS_2_^n^-US^+^ early and late neural pattern similarity were subtracted from CS_1_^−^-US^−^/CS_2_^−^-US^−^ and CS_1_^+^-US^+^/CS_2_^+^-US^+^ early and late neural pattern similarity, respectively. We hypothesized that neural pattern similarity would show changes from early (first 25) to late (last 25) trials, as both CS_2_^+^ and CS_2_^−^ should become more similar to their respective indirectly associated US, indicating associative learning transfer from CS_1_ to CS_2_. Contrarily, CS_1_^+^ and CS_1_^−^ should, if anything, become less similar to the respectively paired US (e.g. due to extinction). Average change of early to late CS-US neural pattern similarity was analyzed at the group level with paired-samples t-tests. Due to the expected negative change for CS_1_^+^ and CS_1_^−^ and the expected positive change for CS_2_^+^ and CS_2_^−^ across trials, we used one-tailed tests accordingly.

### Code Accessibility and Data Availability

Custom analysis code for the reported behavioral data analyses and the multivariate fMRI, behavioral data, extracted classification accuracies, neural pattern similarity matrices and thresholded, unthresholded statistical maps that support the findings of this study are available at GitHub (https://github.com/LLuettgau/higher_order_learning). There are restrictions to the availability of the neuroimaging raw data due to them containing information that could compromise research participant privacy/consent. Neuroimaging raw data are available upon reasonable request from the corresponding author (LL).

## Results

### Behavioral evidence for transfer of motivational value during second-order conditioning

Across two experiments, we found behavioral evidence that our second-order conditioning procedure induced both direct associative learning (CS_1_) and associative transfer learning effects (CS_2_). Our data suggests that second-order conditioning induces preferences for higher-order conditioned stimuli – stimuli that had themselves never been directly paired with a reinforcing event or outcome. Participants (*N* = 49, combined across two experiments) first established Pavlovian visuo-gustatory associations between a visual CS^+^_1_ and an appetitive gustatory US^+^ (chocolate milk or orange juice), and between a visual CS^−^_1_ and an aversive gustatory US^−^ (quinine-HCl) during first-order conditioning (FOC, performed outside MRI, Fig. 1C). During SOC, participants were visually presented with CS_2_^+^ followed by CS_1_^+^, CS_2_^−^ followed by CS_1_^−^, and a pairing of CS_2_^n^ followed by CS_1_^n^ (Fig. 1D). Since CS_1_^n^ had not been presented during first-order conditioning, both CS_2_^n^ and CS_1_^n^ were considered neutral CS, serving as control stimuli in fMRI analyses. Importantly, participants were not informed about the underlying associative structure of the tasks but were instructed to perform simple attentional control tasks. This was aimed at leaving participants unaware of the associative learning process. Post-experimental tests for explicit knowledge indeed revealed that participants were unaware of the (indirect) associations between CS_2_ and US (Table 1). The number of participants that had indeed realized the indirect associations between CS_2_ and US was significantly below the value that would be expected under a random guessing assumption (29%, *P* = 0.004, binomial test vs. 0.5). Participants could also not reliably indicate which CS_2_ and which US had been indirectly linked in the experiment (mean correct responses: *M* = 0.31, *SD* = 0.68, maximum of 2 correct answers possible, Table 1). In a subsequent choice preference test (Fig. 1E), participants performed binary decisions between pairs of CS_1_ and CS_2_. Choice trials were interspersed with lure decisions between CS_1_^n^ or CS_2_^n^ and other lure fractals or kanjis, respectively, that had only been seen during pre-task rating. We reasoned that choice probabilities should exceed the indifference criterion (choice probability > 0.5) if associative direct (CS_1_) and associative transfer (CS_2_) learning effects occurred. Additionally, we assumed that choice probability should not exceed the indifference criterion in trials involving CS^n^-to-lure stimuli, since both CS_2_^n^ and CS_1_^n^ had not been paired with a US and should therefore not have acquired motivational value. Combined across the two experiments, participants showed a preference both for the appetitive first- and second order stimuli, CS_1_^+^ and CS_2_^+^, respectively, over the aversive CS_1_^−^ and CS_2_^−^ (Fig. 1F). Among six candidate Bayesian multilevel generalized linear models, a model that combined individually varying intercepts for each participant with covarying CS-specific intercepts (Equation/Model 5) best captured the observed pattern of choice behavior using Pareto-Smoothed Importance Sampling (PSIS) values (± standard error): Model 5 = 1788.6 (±53.13), Model 6 = 2034.8 (±33.53), Model 4 = 3173.2 (±18.37), Model 2 = 3184.5 (±15.64), Model 3 = 3236.6 (±8.34), Model 1 = 3239.1 (±8.12). We found converging evidence using Widely Applicable Information Criterion (WAIC) values (± standard error): Model 5 = 1782.5 (±52.65), Model 6 = 2033.1 (±33.50), Model 4 = 3173.0 (±18.36), Model 2 = 3184.4 (±15.62), Model 3 = 3236.6 (±8.36), Model 1 = 3239.0 (±8.08). Model 5 had higher predictive accuracy than any other model considered and captured the choice behavior more accurately than the more complex model combining individually varying intercepts, study-specific intercepts with covarying CS-specific intercepts (Equation/Model 6). This suggest that adding information about study (behavioral or fMRI) does not further increase the model’s predictive accuracy - indicating that choice behavior did not differ between the two studies. Within the varying intercepts model (Equation/Model 5), the highest posterior density intervals (HPDI) around the group-level intercept parameter (i.e. the mean posterior choice probability, *μα* .71) for CS_1_^+^ versus CS_1_^−^ [.61; .81] and for CS_2_^+^ versus CS_2_^−^ (*μα* .69) [.56; .81] did not overlap (0% overlap) with the defined ROPE (choice probability = [.45; .55]). Importantly, there was no evidence for both choice probabilities of CS_1_^n^ (*μα* .50) and CS_2_^n^ (*μα* .42) being different from chance level in CS^n^-to-lure comparison choice trials (Fig. 1F, HPDIs: [.38; .62] and [.31; .53], 57.43% and 28.05% ROPE overlap, respectively). This indicates that our conditioning procedure reliably induced both direct associative (CS_1_) and associative transfer learning effects (CS_2_). Together, these data indicate that our conditioning procedures induced both a preference of CS_1_^+^ over CS_1_^−^, and of CS_2_^+^ over CS_2_^−^.

**Table 1.** Explicit knowledge of CS-US associations.

| **CS<sub>1</sub>–US**<br>**(*N* = yes)** | **CS<sub>1</sub>–US**<br>**(*Mean* + *SD*)** | **CS<sub>2</sub>–US**<br>**(*N* = yes)** | **CS<sub>2</sub>–US**<br>**(*Mean* + *SD*)** |
| --- | --- | --- | --- |
| 38 (78 %) | .98 (.95) | 14 (29 %) | .31 (.68) |

*Note:* Number (*N*) and percentage (in parentheses) of participants indicating explicit knowledge of CS–US associations between directly paired CS<sub>1</sub> and US (providing an answer to the question which CS<sub>1</sub> was associated with which US, regardless of being correct or incorrect) and indirectly paired CS<sub>2</sub> and US (responding “yes” to the question whether there was a relationship between CS<sub>2</sub> and US). These numbers of participants were significantly above (CS<sub>1</sub>–US) and below (CS<sub>2</sub>–US) the numbers expected under the assumption of random guessing, $P < .001$ and $P = .004$ , respectively (Binomial test vs. 0.5). Additionally, we provide the mean (and standard deviation, *SD*) number of correct answers for CS–US associations between CS<sub>1</sub> and US, and CS<sub>2</sub> and US. Since participants were presented with two CS–US associations during conditioning, the maximum number of correct answers is 2 in both cases.

### CS and US ratings

#### CS_1_ ratings

During the ratings prior to the learning experiment, the subjective values/liking for the three fractals selected as CS_1_ were higher for CS_1_^n^ than CS_1_^−^ (*t*_48_ = 2.26, *P* = .028, paired-samples t-test). Of note, this difference was only significant within the fMRI study (CS_1_^n^ > CS_1_^−^ : *t*_28_ = 2.44, *P* = .022, paired-samples t-test). We reason that the observed difference most likely resulted from the selection process and the inherent rank-ordering of the CS according to subjective value ratings. However, there were no significant differences between CS_1_^n^ and CS_1_^+^ (*t*_48_ = 1.68, *P* = .101, paired-samples t-test). Most importantly, no rating differences were observed between CS_1_^−^ and CS_1_^+^ (*t*_48_ = 0.44, *P* = .659, paired-samples t-test), indicating that there were no systematic (and unintended) differences between the selected fractals that were later paired with US^−^ or US^+^. A rmANOVA evaluating changes in pre-post ratings indicated that none of the CS_1_ was rated significantly different from the others across ratings (main effect of stimulus: *F*_2,96_ = 3.09, *P* = .050, η^2^*_p_* = .06, rmANOVA). There was also no significant pre-post change in ratings across all CS_1_ (main effect of time: *F*_1,48_ = 2.02, *P* = .161, η^2^*_p_* = .04, rmANOVA). However, there was a significant CS x time interaction effect (*F*_2,96_ = 3.36, *P* = .039, η^2^*_p_* = .07, rmANOVA), that was driven by more positive change of CS_1_^+^ ratings compared to CS_1_^−^ (*t*_48_ = 2.06, *P* = .045, paired-samples t-test on pre-post differences) and CS_1_^n^ (*t*_48_ = 2.52, *P* = .015, paired-samples t-test on pre-post differences), but no significant difference between CS_1_^−^ and CS_1_^n^ (*t*_48_ < .01, *P* > .999, paired-samples t-test on pre-post differences). The CS x time interaction effect was marginally significant in the behavioral study (*F*_2,38_ = 2.78, *P* = .075, η^2^*_p_* = .13, rmANOVA), but was not significant in the fMRI study (*F*_2,56_ = 1.17, *P* = .318, η^2^*_p_* = .04, rmANOVA). In the behavioral study, there was more positive change of CS_1_^+^ ratings compared to CS_1_^n^ (*t*_19_ = 2.97, *P* = .008, paired-samples t-test on pre-post differences), but all pairwise comparisons were not significant within the fMRI study (all *P*s > .184).

#### CS_2_ ratings

None of the three kanjis selected as CS_2_ differed significantly from each other during the pre-experimental rating (all *P*s > .919), indicating that there were no systematic (and unintended) differences between the selected kanjis that were later paired with CS_1_^−^ or CS_1_^+^. Similarly, in an rmANOVA evaluating changes in pre-post ratings, none of the CS_2_ was rated significantly different from the others across both ratings (main effect of stimulus: *F*_2,96_ = .46, *P* = .630, η^2^*_p_* = .01, rmANOVA). There was significant pre-post change in ratings across all CS_2_ (main effect of time: *F*_1,48_ = 4.11, *P* = .048, η^2^*_p_* = .08, rmANOVA), resulting from slightly overall higher ratings after learning (mean difference: *M* = .29). This effect was marginally significant in the behavioral study (*F*_2,38_ = 3.34, *P* = .083, η^2^*_p_* = .149, rmANOVA), but was not significant in the fMRI study (*F*_2,56_ = 1.31, *P* = .263, η^2^*_p_* = .05, rmANOVA). However, there was no significant CS x time interaction effect for CS_2_ ratings (*F*_2,96_ = .89, *P* = .416, η^2^*_p_* = .02, rmANOVA).

Together, both CS_1_ and CS_2_ rating results indicate that the conditioning procedure only induced reliable changes in subjective value/liking of CS_1_^+^ (compared with CS_1_^n^) in the behavioral study, but these changes were not found in the fMRI study. CS_2_ ratings did not exhibit systematic changes in subjective value/liking.

#### US ratings (learning experiment)

Overall, US valence ratings did not differ from pre to post rating across both US (main effect of time (pre/post): *F*_1,48_ = 0.34, *P* = .564, η^2^*_p_* = .01, rmANOVA), but the appetitive US^+^ was rated as significantly more pleasant than the aversive US^−^ (main effect of stimulus: *F*_1,48_ = 466.64, *P* < .001, η^2^*_p_* = .91, rmANOVA). There was also a significant interaction effect between stimulus and time, indicating that US^+^ was rated as more pleasant and US^−^ was rated as less pleasant from pre to post (interaction effect of stimulus x time: *F*_1,48_ = 7.32 *P* = .009, η^2^*_p_* = .13, rmANOVA). Importantly, no subject rated US^−^ as more pleasant than US^+^ (pre or post). These results indicate that both US^−^ and US^+^ possessed the intended reinforcing properties and valence, both before and after being used as unconditioned stimuli in first-order conditioning. Additionally, these reinforcing properties were even enhanced over the course of conditioning, ruling out potential exposure-dependent habituation or devaluation of the reinforcers. In the fMRI study, intensity ratings did not differ between US overall (main effect of stimulus: *F*_1,28_ = 1.05, *P* = .315, η^2^*_p_* = .04, rmANOVA), but US were rated as significantly more intense at the post rating (main effect of time: *F*_1,28_ = 4.89, *P* = .035, η^2^*_p_* = .15, rmANOVA). There was also a significant interaction effect between stimulus and time, indicating that US^−^ intensity rating increased more strongly than US^+^ from pre to post (interaction effect of stimulus x time: *F*_1,28_ = 5.62, *P* = .025, η^2^*_p_* = .17, rmANOVA). The average temperature (°Celsius) of US^+^ (*M* = 22.49, *SD* = 1.46) was significantly higher than temperature of US^−^ (*M* = 22.37, *SD* = 1.52) before being loaded into the syringes (*t*_27_ = 2.22, *P* = .035, paired-samples t-test). However, we would argue that this minor difference in average temperatures (*M* = .12) is unlikely to be perceived by participants. Additionally, this difference does not meaningfully influence the observed results, since no neural activation was recorded during first-order conditioning and the multivariate classification analysis was trained on the data resulting from the fMRI classifier training experiment.

#### US ratings (classifier training experiment)

In the fMRI classifier training experiment, overall US valence ratings were higher in pre than in post rating (main effect of time: *F*_1,28_ = 11.67, *P* = .002, η^2^*_p_* = .29, rmANOVA), but the appetitive US^+^ was rated as significantly more pleasant than the aversive US^−^ across ratings (main effect of stimulus: *F*_1,28_ = 185.28, *P* < .001, η^2^*_p_* = .87, rmANOVA). There was no significant interaction effect between stimulus and time (*F*_1,28_ = 0.28, *P* = .602, η^2^*_p_* = .01, rmANOVA). Importantly, no subject rated US^−^ as more pleasant than US^+^ (pre or post). Intensity ratings for US^+^ were higher than for US^−^ across ratings (main effect of stimulus: *F*_1,28_ = 15.14, *P* < .001, η^2^*_p_* = .35, rmANOVA). There was no difference between US ratings from pre to post (main effect of time: *F*_1,28_ = 0.02, *P* = .899, η^2^*_p_* < .001, rmANOVA). However, there was also a marginal interaction effect between stimulus and time, indicating that US^−^ intensity rating increased while US^+^ intensity rating decreased from pre to post (interaction effect of stimulus x time: *F*_1,28_ = 3.93, *P* = .057, η^2^*_p_* = .12, rmANOVA). Average temperature of US^+^ (*M* = 21.71, *SD* = 1.93 °C) and US^−^ (*M* = 21.68, *SD* = 1.95) did not differ before being loaded into the syringes (*t*_27_ = 0.71, *P* = .486, paired-samples t-test).

### First-order CS reinstate neural US patterns during second-order conditioning

After establishing behavioral evidence for associative transfer learning, we reasoned that a prerequisite for the observed learning effect would be reinstatement of the neural US pattern by the paired CS_1_ to establish a direct associative link between CS_2_ and US. Importantly, it should be noted that US reinstatement during SOC could only be faithfully attributed to the respective CS_1_, but not to CS_2_, since only CS_1_ had been directly paired with the US, and CS_2_ had not previously been experienced. To test for cortical reinstatement of neural patterns representing US during second-order conditioning, we used functional magnetic resonance imaging (fMRI) and a cross-session (Stokes et al. 2009), cross-modality searchlight (Kriegeskorte et al. 2006) classification approach (multivariate pattern analysis, MVPA). To obtain unbiased estimates of the neural patterns representing our gustatory US, without the confounding influence of associations to a learned first-order CS, we first performed an fMRI classifier training experiment on day 1 (Fig. 1B) during which the US^+^ and US^−^ were presented. A multivariate pattern classifier (linear support vector machine, C-SVM) was trained on the fMRI data from this session.

On day 2, during SOC, we used the weights of the classifier trained on gustatory neural patterns to predict the class label of the visual CS (Fig. 1D) that had been paired with the US during FOC (Fig. 1C, see Fig. 3A for schematic of the classification approach). In other words, we used data from one sensory modality, assessed during a first day to train a classification model and tested the model’s generalizability to unseen data from another sensory modality on a second day. We found evidence for reinstatement of US patterns in a region in the left lateral orbitofrontal cortex (lOFC). In this region, it was possible to predict the class labels of neural CS patterns based on US pattern information obtained during classifier training on the previous day. Classification with a 3-mm searchlight revealed a small-volume corrected cluster of above-chance level (0.5) classification accuracy (extracted cluster mean = 0.56, *SD* = 0.07) in the left lateral OFC (lOFC, Fig. 3B, peak voxel at MNI [x = –21, y = 30, z = –17], *z* = 2.26, *P* = 0.012, random-effect cluster corrected (Stelzer et al. 2013), 50,000 iterations, one-tailed). The location of this lOFC cluster is consistent with this region’s well-documented role in gustatory processing, particularly in representing motivational (Small et al. 1999; Rolls 2000, 2006) aspects of gustatory sensation and taste memory (Kobayashi et al. 2004). To ensure that our results are not dependent on this particular choice of ROI, we repeated the same analysis using two different, independent ROIs. The first was an anatomical mask of lateral orbitofrontal cortex, the second was obtained from an independent gustatory mapping study by Benz and colleagues (K. Benz, personal communication, 12/2019). We found similar results in the left lOFC anatomical ROI (peak voxel at MNI [x = –21, y = 30, z = –17], *z* = 1.96, *P* = .024, corrected, one-tailed) and in the mask from Benz and colleagues (peak voxel at MNI [x = –21, y = 30, z = –17], *z* = 1.84, *P* = .033, corrected, one-tailed).

**Figure 2.**
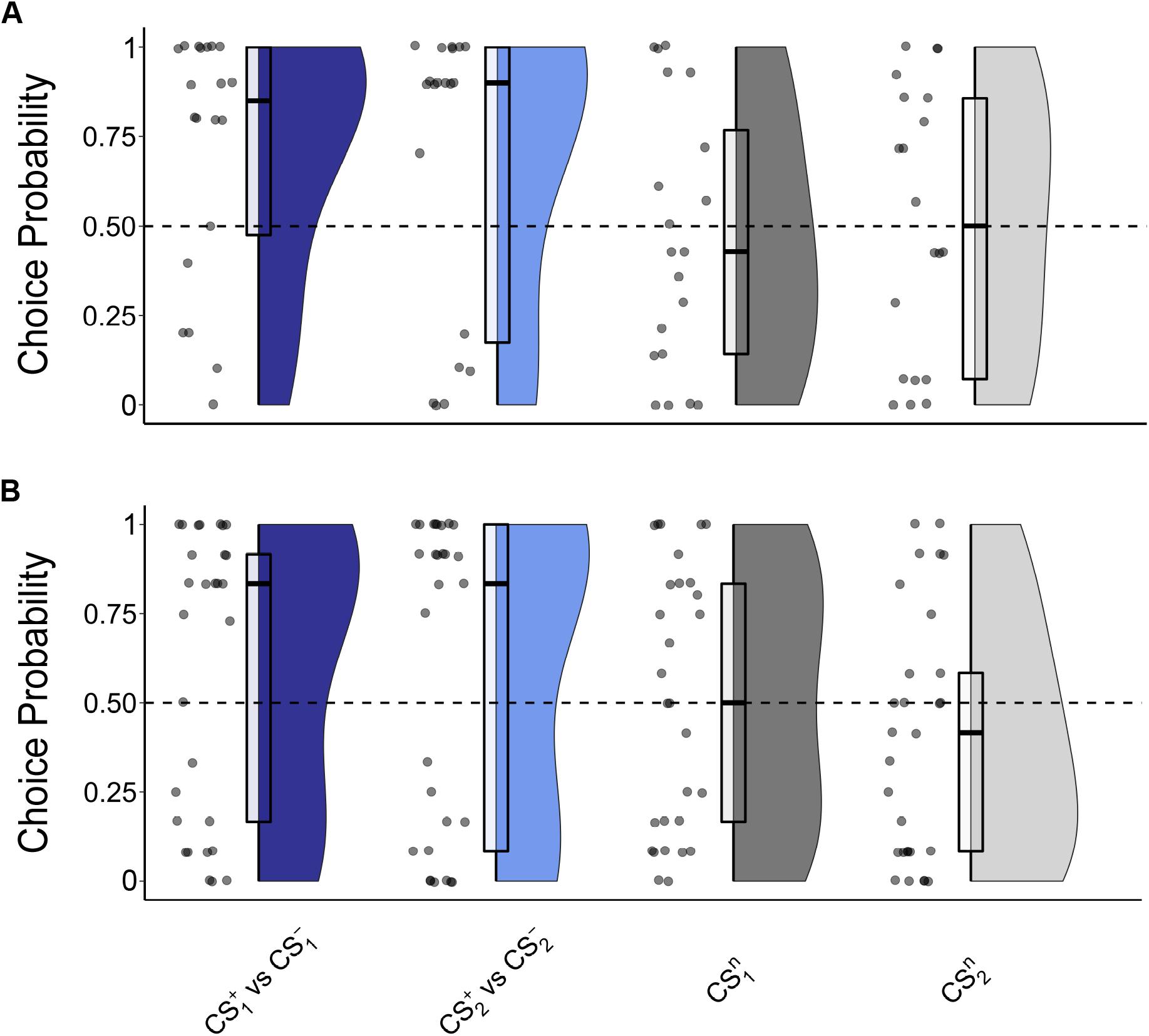
Behavioral results separated for both studies. Results as in 1F, but separately for the behavioral (A, *N* = 20) and fMRI study (B, *N* = 29). Raincloud plots showing density of choice probability. Box plot center lines represent (pre-averaged) study medians and box bottom/top edges show 25^th^/75^th^ percentile of the (pre-averaged) data, respectively.

**Figure 3.**
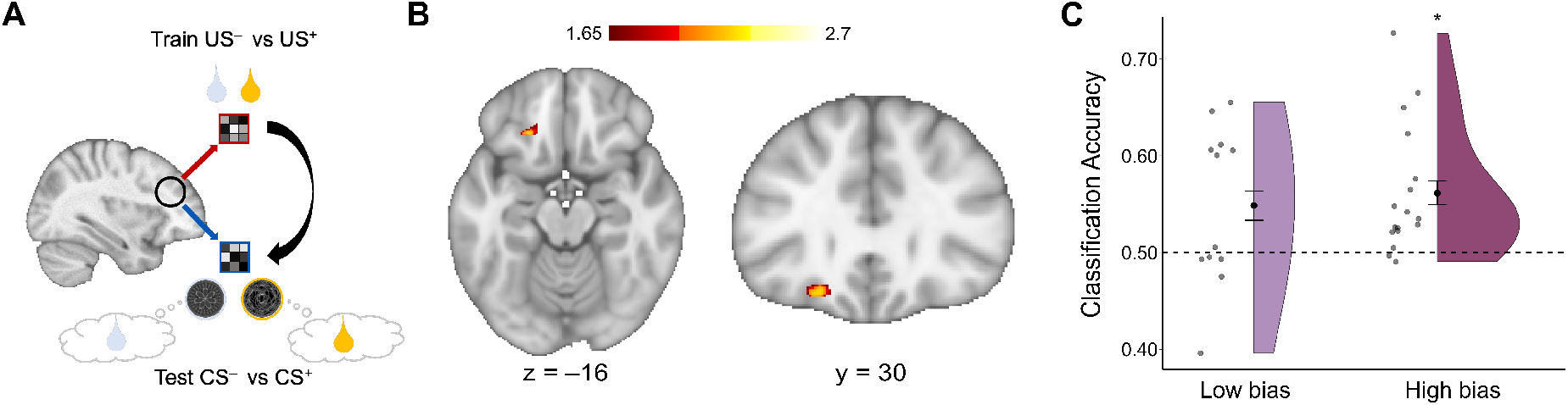
Reinstatement of US representations during second-order conditioning. A) Analysis approach: For cross-session, cross-modality classification, a multivariate classifier was trained on the spatial activation patterns (*t* maps) of US^+^ versus US^−^ (obtained from the classifier training session) and tested on CS_2_^+^/CS_1_^+^ versus CS_2_^−^/CS_1_^−^ during SOC. CS_2_^n^/CS_1_^n^ related activations were subtracted to control for general visual effects. B) Classification accuracy exceeded chance level (0.5) in a small-volume corrected cluster in the left lateral orbitofrontal cortex (lOFC), using random-effect cluster-statistics (Stelzer et al. 2013). C) Average classification accuracy in the lOFC cluster (extracted cluster mean = 0.56, *SD* = 0.07) was significantly above chance level in the high bias group of participants showing higher than chance preference for CS_2_^+^, but not in the low bias group. However, there was no significant difference in classification accuracies in lOFC between the high bias and low bias group. Color bar represents *Z*-values. Black dots indicate means and error bars represent standard errors of the means of the data, respectively.

Very similar classification results could be obtained in the left lOFC if neural activation related to CS_n_ was not subtracted from both CS_2_^+^/CS_1_^+^ versus CS_2_^−^/CS_1_^−^ patterns before classification (as reported in the previous analysis).

When we split participants into a high- and low-bias group, depending on their preference for the CS_2_^+^, we observed that significant classification of CS labels in lOFC was possible in the high bias group (*Z* = 3.43, *P* = .006, *U3_1_* = .88, one-sample Wilcoxon signed-rank test, two-tailed), but not in the low bias group (*Z* = 1.44, *P* = .151, *U3_1_* = .58, one-sample Wilcoxon signed-rank test, two-tailed, Fig. 3C). However, there was no significant difference in classification accuracies in lOFC between the high bias and low bias group (*Z* = 0.64, *P* = .260, *U3* = .50, Mann-Whitney U test, one-tailed). In an additional exploratory analysis testing for sex differences, we found that classification accuracies in the lOFC did not differ between female (*Median* = .53) and male (*Median* = .55) participants (*Z* = 0.50, *P* = .616, *U3* = .64, Mann-Whitney U test).

### Interaction between lOFC, amygdala, and medial OFC during second-order conditioning

To form an associative link between CS_2_ and US, the reinstated US patterns need to be projected from their cortical storage site to regions like amygdala and hippocampus, allowing for convergence of US and CS_2_ information. We investigated BOLD signal covariation of the cluster in the left lOFC with the whole brain during SOC using two separate psychophysiological interaction (PPI) analyses. In the first PPI, we investigated general covariation differences in response to CS^−^ and CS^+^ relative to CS^n^, reasoning that in the former, but not in the latter, reinstated US representations need to be linked with the CS. We found that covariation of BOLD signal in the left lOFC with a cluster in the left hippocampus, extending to amygdala and medial temporal lobe (Fig. 4A, peak voxel at MNI [x = –38, y = –11, z = –24], Z = 3.75, *P* = .003, whole-brain corrected) and a cluster in the right inferior temporal gyrus, extending to the right hippocampus (peak voxel at MNI [x = 40, y = –32, z = –20], Z = 3.55, *P* = 0.007, whole-brain corrected) was higher in CS^−^ and CS^+^ compared to CS^n^ trials. Despite the fact that we used an approach with high statistical sensitivity for group-level cluster-based inference (FSL’s FLAME1+2 (Beckmann et al. 2003)), a robust estimation technique that yields low family-wise errors rates even at liberal activation thresholds (*Z* > 2.3, corresponding to *p* < 0.05) (Eklund et al. 2016), we would like to note that the aforementioned result did not survive whole-brain correction at more conservative thresholds (e.g. *Z* > 2.6). Additionally, it should be noted that the observed BOLD covariation differences could potentially also be explained by higher familiarity and salience of CS^−^ and CS^+^ relative to CS^n^ and cannot be attributed exclusively by the formation of associative links with a US. Furthermore, in the same PPI analysis, we found a positive correlation between second-order choice preference and functional covariation of the left lOFC with a region in the right and left lateral prefrontal cortex (Fig. 5, right peak voxel at MNI [x = 49, y = 41, z = 7], Z = 3.80, *P* < 0.001, whole-brain corrected; left peak voxel at MNI [x = –45, y = 46, z = –4], Z = 3.93, *P* = 0.010, whole-brain corrected). The more participants preferred CS_2_^+^ (versus CS_2_^−^), the stronger these regions’ BOLD signal covaried during SOC. Some studies hint at complex interactions between OFC and amygdala that differ between appetitive and aversive stimuli (Morrison et al. 2011). Therefore, we next asked whether there were any specific differences between CS^−^ and CS^+^ trials in lOFC BOLD signal covariation with other regions. We found that covariation of the BOLD signal in the left lOFC was higher in CS^+^ compared to CS^−^ trials in a cluster located in the medial OFC, extending to subgenual anterior cingulate cortex (Fig. 4B, peak voxel at MNI [x = 10, y = 50, z = –1], Z = 3.90, *P* < 0.001, whole-brain corrected) and a cluster in the left anterior insula, extending to left caudo-lateral OFC and temporal pole (peak voxel at MNI [x = –51, y = 18, z = –11], Z = 3.33, *P* = 0.004, whole-brain corrected). Given both the difficulties in inferring directionality from fMRI data and the reciprocal nature of connections between lOFC and both amygdala and medial OFC, a directionality of our PPI results cannot be determined. Again, exploratory analyses testing for sex differences revealed no significant differences in either the PPI analysis comparing CS^−^/CS^+^ vs CS^n^ (parameter estimates extracted from amygdala mask) or in the PPI analysis comparing CS^−^ vs CS^+^ trials (parameter estimates extracted from mOFC mask) between female and male participants (*Z* = 1.07, *P* = .285, *U3* = .64, Mann-Whitney U test and *Z* = .98, *P* = .326, *U3* = .57, Mann-Whitney U test, respectively).

**Figure 4.**
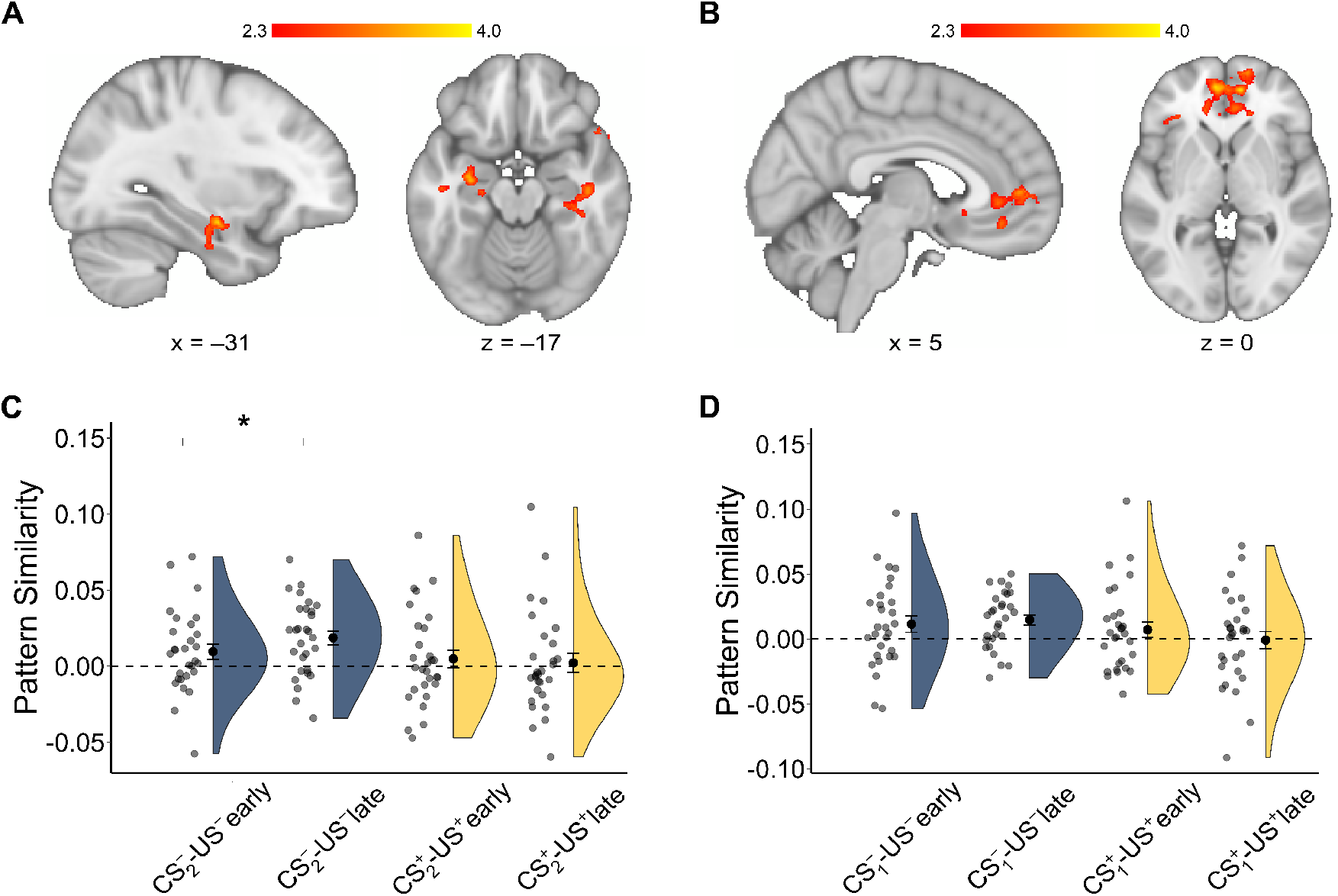
Psychophysiological interaction analyses and neural pattern similarity results. A) BOLD covariation was higher in CS^−^ and CS^+^ trials than in CS^n^ trials between the left lOFC and a cluster in the anterior hippocampus, extending to (basolateral) amygdala and medial temporal lobe and a cluster in the right inferior temporal gyrus, extending to the right hippocampus. B) Covariation of BOLD signal in the left lOFC was higher in CS^+^ trials compared to CS^−^ trials in a cluster located in the medial OFC, extending to subgenual anterior cingulate cortex. C, D) Template-based neural pattern similarity between US patterns from the classifier training experiment and CS_2_ (C) and CS_1_ (D) patterns during SOC, separately estimated for early (first 25 trials) and late (last 25 trials) of SOC in a bilateral amygdala ROI. C) CS_2_^−^ and US^−^ patterns became more similar from early to late trials. However, there was no evidence for a difference between early and late trial similarity for CS_2_^+^ and US^+^. D) There was no evidence for change in similarity for CS_1_^−^-US^−^ or CS_1_^+^-US^+^ neural pattern similarity. Black dots indicate means and error bars represent standard errors of the means of the data, respectively. Color bars represent *Z*-values.

**Figure 5.**
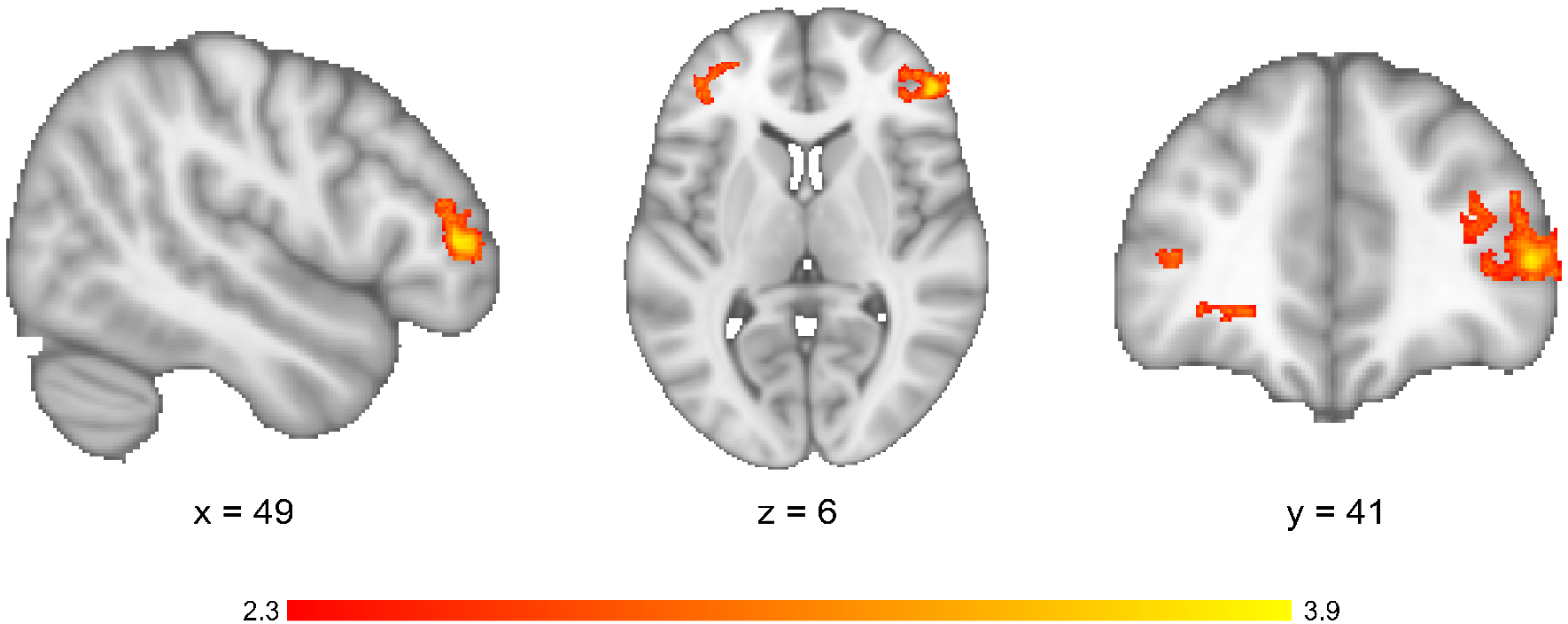
Psychophysiological interaction analyses: Association with second-order choice preferences. A positive correlation was observed between second-order choice preferences (CS_2_^+^ versus CS_2_^−^) and functional covariation between the left lOFC and a region in the right and left lateral prefrontal cortex. The more participants preferred CS_2_^+^ (versus CS_2_^−^), the higher these regions’ activation covaried during second-order conditioning. Color bar represents *Z*-values.

### Plasticity of association between second-order CS and US in the amygdala

We reasoned that if the amygdala uses the reinstated cortical outcome pattern in lOFC to acquire an association between CS_2_ and the respective US, one would expect similarity between the neural patterns evoked by CS_2_ and US, respectively. This similarity should increase over the course of second-order conditioning, as the association is being acquired. Importantly, during the classifier training experiment, there was evidence for US representations in the amygdala. We found differential neural patterns in response to US^−^ and US^+^ in the amygdala (among other regions) using an exploratory leave-one-run-out cross-validated classification analysis on the neural pattern recorded during the classifier training experiment (Fig. 6). Using a least-squares separate (LS-S) approach (Mumford et al. 2012) to deconvolve single-trial estimates, we first computed overall neural pattern similarity (Kriegeskorte et al. 2008) between the pattern evoked by CS_2_ (during SOC) and their respective US (during classifier training) in the bilateral amygdala. There was significant pattern similarity between CS_2_^−^ and US^−^ (*t*_28_ = 3.38, *P* = 0.002, *U3_1_* = 0.79; but not between CS_2_^+^ and US^+^, *t*_28_ = 0.74, *P* = 0.464, *U3_1_*= 0.55). The reasons for this difference between CS_2_ are unknown, but their investigation might present a fruitful avenue for future studies. Due to the observed disparity, we reason that it is possible that the observed preference for CS_1_^+^ and CS_2_^+^ was indeed due to an aversion against CS_1_^−^ and CS_2_^−^. If the observed neural pattern similarity between CS_2_ and US indeed reflects the formation of an associative link during second-order learning, one would expect the observed similarity to increase from early to late stages of learning, as the association is being acquired and the CS_2_ comes to predict the US. We therefore compared our measure of neural pattern similarity between early and late trials of SOC. As expected, neural pattern similarity between CS_2_^−^ and US^−^ in the bilateral amygdala ROI increased from early to late trials of SOC (*t*_28_ = 1.88, *P* = 0.035, Cohen’s *d* = 0.35, paired-samples t-test, one-tailed, Fig. 4C). This change in similarity was not observed for the first-order CS_1_^+^-US^+^ or CS_1_^−^-US^−^ pattern similarity (Fig. 4D), nor for second-order CS_2_^+^-US^+^ similarity (all *P* > .310, paired-samples t-tests, one-tailed, Fig. 4C).

**Figure 6.**
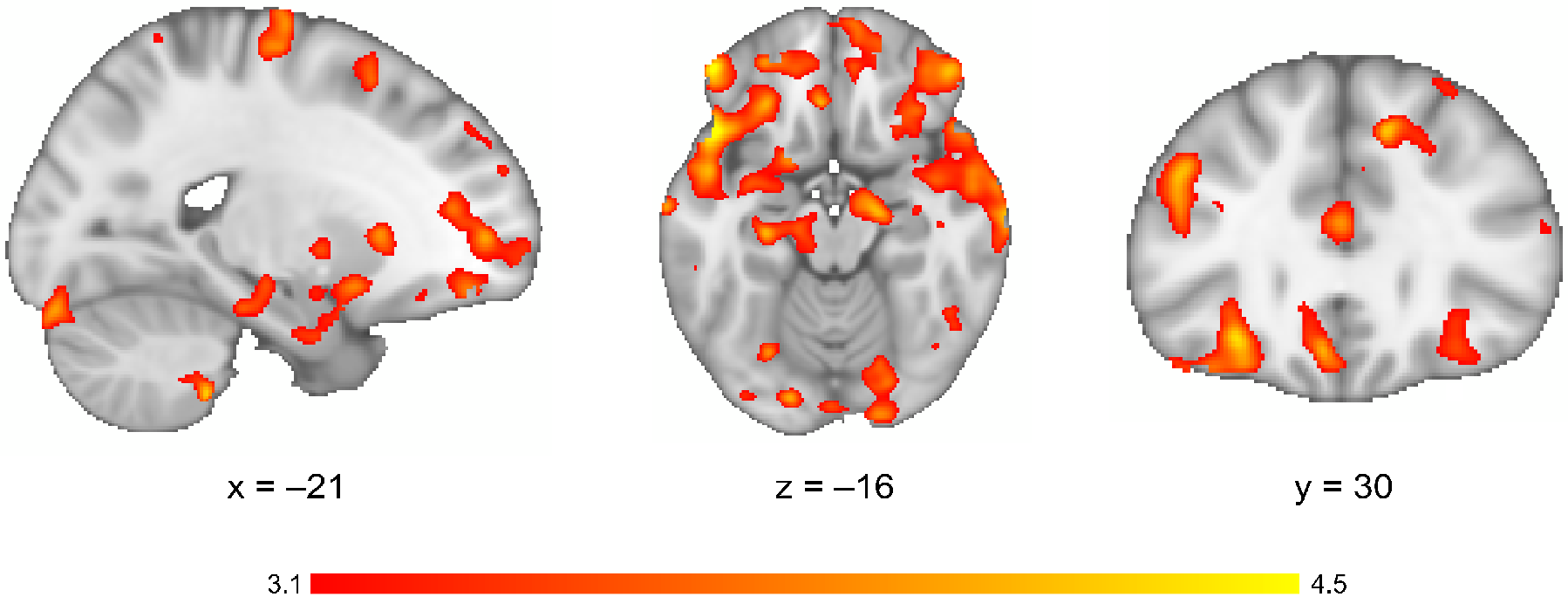
US^+^ versus US^−^ classification. Group average (*N* = 29) cross-validated predictive accuracy for US^+^ versus US^−^ activation patterns during the classifier training experiment. As an exploratory analysis, we used multivariate searchlight classification (3-mm searchlight spheres) to distinguish between the spatial activation patterns (*t* maps) of US^+^ versus US^−^ across the whole brain, resulting in one whole-brain map of classification accuracies per participant. This classification was performed using a 5-fold cross-validation scheme. Data was averaged across participants and smoothed with a Gaussian kernel with 5 mm FWHM. Please note that due to the exploratory nature and illustrative purpose of this analysis, no multiple comparisons correction using random-effects cluster-statistics was performed. Color bar represents uncorrected *Z*-values.

## Discussion

Using a second-order conditioning paradigm, we provided evidence for associative transfer of value during higher-order conditioning. Decisions were biased by values indirectly acquired by second-order learning. Participants were more likely to select directly and indirectly appetitively paired stimuli over aversively paired stimuli, closely resembling rodent studies describing instrumental behavior guided by second-order conditioning (Sharpe et al. 2017; Maes et al. 2020). Notably, choice biases for second-order conditioned stimuli in the present study emerged in the absence of explicit knowledge of the underlying higher-order associative structure. This suggests that humans, similar to rodents, implicitly acquire preferences through higher-order transfer learning mechanisms – extending previous studies promoting acquisition of explicit associative relationships between stimuli (Jara et al. 2006; Pauli et al. 2019; Wang et al. 2020).

The present study thus demonstrates that human value-based decision making is affected by motivational value implicitly conferred via second-order conditioning. Our study is – to the best of our knowledge – the only report so far demonstrating that human binary decision making is governed by motivational value transfer via second-order conditioning. Although there is a rich literature on second-order conditioning, direct evidence for this phenomenon in humans using similar procedures as in previous animal work was lacking. Our study enables a closer comparison between humans and other species during higher-order learning. Moreover, even animal studies, albeit well-covering Pavlovian settings (Gewirtz and Davis 2000; Sharpe et al. 2017), have thus far not reported effects of second-order conditioning on binary choice behavior.

Thus far, most studies investigating higher-order learning effects in humans have used sensory preconditioning procedures (Wimmer and Shohamy 2012; Kurth-Nelson et al. 2015; Wang et al. 2020). Theoretical accounts and empirical findings suggest that the value structure acquired during sensory preconditioning and second-order conditioning – despite the close conceptual links between paradigms – differs fundamentally. In sensory preconditioning, a stimulus-stimulus association is acquired first (preconditioning phase), before one stimulus is paired with an outcome (conditioning phase). It has been found that preconditioned stimuli do not directly acquire value, as they do not serve as conditioned reinforcers (Sharpe et al. 2017). Instead, preconditioned stimuli reactivate both conditioned stimuli and indirectly associated outcome representations at preference assessment (Barron et al. 2020; Wang et al. 2020), suggesting value inference based on associative chaining. Contrarily, during second-order conditioning, second-order CS directly acquire value, allowing them to act as conditioned reinforcers (Sharpe et al. 2017). These findings indicate that associative transfer of motivational value from outcomes to higher-order, indirectly paired CS are observed in second-order conditioning, but not in sensory preconditioning.

We found reinstatement of neural representations of gustatory outcomes by directly paired visual first-order CS. This reinstatement occurred in a region in the rostrolateral OFC, which has previously been implicated in representing stimulus-outcome associations (Klein-Flügge et al. 2013; Jocham et al. 2016; Luettgau et al. 2020) and in correctly assigning credit for a reward to the causal stimulus choice (Walton et al. 2010; Jocham et al. 2016). It is also part of the network most consistently involved in taste processing (Neurosynth, Yarkoni et al., 2011) and plays a well-documented role in representing motivational aspects of gustatory sensation (Small et al. 1999; Rolls 2000, 2006) and taste memory (Kobayashi et al. 2004). Importantly, we had trained the classifier on gustatory US prior to pairing them with first-order CS (on a separate day). Thus, decoding of the visual CS during second-order learning cannot be spuriously driven by visual responses elicited by the US or the visual swallowing cue during classifier training. Despite the fact that the classifier was tested on the entire activation pattern elicited by the CS_1_-CS_2_-pairs, US reinstatement during second-order conditioning is most likely attributable to the respective CS_1_. This is due to the fact that, despite the close temporal proximity with CS_2_, only CS_1_, but not CS_2_, had been directly paired with the outcome and should thus elicit a US representation (Wagner 1981). Since we had optimized our design for behavioral second-order learning effects by introducing only small temporal offsets between first- and second-order CS, methods with higher temporal resolution than fMRI would be required to dissociate the specific responses to CS_1_, and CS_2_.

OFC clearly does not act in isolation. The OFC region in which we observed reinstatement of outcome representations displayed increased task-related BOLD signal covariation with amygdala, anterior hippocampus, and medial OFC during second-order learning. Neurons in OFC and amygdala show complex, bi-directional interactions during acquisition and reversal of outcome-predictive associations (Morrison et al. 2011). Functional disconnection of OFC and amygdala using asymmetric lesions produces deficits in flexibly adjusting behavior to changes in stimulus value (Baxter et al. 2000; Fiuzat et al. 2017). We also observed that representations of visual second-order CS in amygdala became more similar to gustatory US representations from early to late phases of second-order conditioning. This indicates development of an associative link between CS_2_ and US, consistent with the finding that both amygdala and hippocampal lesions impair second-order learning (Gewirtz and Davis 1997; Gilboa et al. 2014).

Similar to the present results, two previous fMRI studies in humans used sequential Pavlovian conditioning paradigms to investigate higher-order learning. Whereas Seymour and colleagues (Seymour et al. 2004) elegantly show that human learning about sequentially presented pain-predictive stimuli relies on the formation of stimulus-stimulus associations and follows temporal difference learning algorithms, putatively implemented in the striatum, Pauli and colleagues (Pauli et al. 2019) provide evidence for model-based representation of stimulus-stimulus associations in the lateral OFC. However, in both paradigms, two CS and a US are presented in close temporal proximity within the same trial. This approach differs from the present design in two important ways. Firstly, unlike second-order conditioning, the paradigm employed by Pauli and colleagues does not require any reinstatement of US representations, nor any transfer of value, but rather imposes different temporal offsets between CS and US. This is in contrast to second-order conditioning, where CS_2_ is presented in a separate block that is entirely devoid of US presentations. Secondly, in both studies, it is difficult to dissociate the activation patterns related to the proximal CS (Pauli et al. 2019) or Cue B/D (Seymour et al. 2004) (CS_1_ in second-order conditioning), CS-related anticipatory pre-activation of US representations and the neural activation related to the US presentation. In contrast, here we trained the classifier on the (gustatory) US on a separate day and then tested on the (visual) CS presented during the second-order learning phase.

Another fMRI study in humans using multivariate pattern classification provides evidence that – similar to the present study – neural pattern similarity between reward-predictive cues and rewards increases over the course of Pavlovian conditioning. This indicates that, as the association is learned, the reward-predictive cue comes to elicit the neuronal representation of the reward (Kahnt et al. 2011). The current study extends on these findings, firstly by showing that first-order stimuli are capable of reinstating neural outcome representations even in a later learning stage (second-order learning), in the complete absence of outcomes, and secondly, by demonstrating that indirectly paired (second-order) stimuli similarly acquire higher-order outcome-predictive properties that guide decision making. Furthermore, in the present study, we used primary reinforcers (as opposed to monetary rewards in Kahnt et al., 2011), which enables a closer comparison across species. We also classified visual stimuli in a phase that was devoid of any direct exposure to reinforcement. Since we trained the classifiers on activation patterns related to gustatory stimuli that had been presented on a separate day, it was possible to mitigate potential biases arising from classification within the same sensory modality and from performing cross-validation within data from the same fMRI session.

Our study presents a neural mechanism for associative transfer learning in second-order conditioning, which thus far has remained unclear. At least four potential mechanisms have been proposed (Rizley and Rescorla 1972; Barnet et al. 1991; Gewirtz and Davis 2000). Three of these suggest that CS_2_ could become associated with a CR using (i) the associative link between stimulus representations of CS_2_ and CS_1_, (ii) direct pairing of the CS_1_-evoked CR with CS_2_, or (iii) because CS_1_ reactivates US representations which then evoke a CR that becomes paired with CS_2_. Previous studies clearly suggest that neither of these three hypotheses can account for second-order learning effects (Rizley and Rescorla 1972; Barnet et al. 1991). It has been shown that CRs towards CS_2_ are unaffected by CS_1_ extinction after second-order conditioning (Rizley and Rescorla 1972; Barnet et al. 1991). Additionally, the magnitudes of CRs for CS_2_ and CS_1_ appear largely uncorrelated (Barnet et al. 1991). Together, these previous findings make it highly implausible that CS_2_–CS_1_ associations are necessary for the expression of CS2-related CRs, or that CS_2_ could be associated with CRs elicited by CS_1_. Here, we provide evidence for a fourth possibility (Gewirtz and Davis 2000; Parkes and Westbrook 2011): that CS_2_ is directly paired with a neural representation of the US, or with the motivational state conveyed by the US. Our results suggest that, during second-order conditioning, outcome representations are reinstated in the lateral OFC. Information of reinstated outcome representations could be communicated between lOFC and amygdala/anterior hippocampus for associative linking between neutral stimuli and outcomes. It should be noted that the present design does not allow us to dissociate whether the reinstated US representations detected by the classification algorithm pertain to the value, intensity or sensory features of the US – or a combination of these features. However, the question which of these features was reinstated by CS_1_ and contributed to the associative transfer of value to CS_2_ is beyond the scope of the present study. Future studies should aim at replicating and maximizing the present, rather subtle higher-order and neural reinstatement effects. This could be achieved by using more strongly aversive (e.g., more aversive gustatory stimuli or mild electric shocks) and appetitive stimuli, presumably combined with even more enhanced periods of food – or even water – deprivation in volunteers.

How the reinstatement of cortical US patterns found in our study relates to the observation in rats that midbrain dopamine neurons acquire temporal difference error signals in response to CS_2_ (Maes et al. 2020) presents an important question for future studies. Furthermore, it would be of great interest to elucidate the directionality and exact content of information flow between lOFC and amygdala/anterior hippocampus, and whether this transfer of information is supported by phase coherence in theta oscillations (Benchenane et al. 2010; Young and Shapiro 2011; Knudsen and Wallis 2020).

Taken together, our data support the idea that during second-order conditioning, second-order CS and representations of outcomes – events that had never been explicitly paired during the individual’s learning history – can be linked by exploiting the relational structure of events. The present study enables a closer comparison between humans and other species during higher-order learning. In conclusion, our results suggest a neural mechanism by which outcome representations can be propagated to stimuli that are never experienced in contiguity with reinforcement, allowing for credit assignment in real-world learning scenarios with infrequent direct encounters with rewards or punishments.

## Acknowledgments

The authors thank all volunteers who participated in this study. The authors would like to thank Halla Mulla-Osman, Stefanie Linnhoff, Leonie Oevel and Denise Scheermann for their invaluable support during data acquisition. Additionally, we thank Gunnar and Christian Lüttgau for electrical engineering support during construction of the gustometer, Karsta Benz for helpful discussions during the set-up of the study and Denise Kramer for production of the gustatory stimuli. We thank the Center for Magnetic Resonance Research of the University of Minnesota for providing the Multiband accelerated fMRI sequence. Computational infrastructure and support were provided by the Centre for Information and Media Technology at Heinrich Heine University Düsseldorf. **Author contributions:** LL designed the study and conceptualized research, acquired the data, analyzed the data, drafted the manuscript, read and edited versions of the manuscript and approved the final version of the manuscript. EP analyzed the data, read and edited versions of the manuscript and approved the final version of the manuscript. CT set up the MRI acquisition protocol, read and edited versions of the manuscript and approved the final version of the manuscript. GJ designed the study and conceptualized research, analyzed the data, read and edited versions of the manuscript and approved the final version of the manuscript.

